# SeqStain using fluorescent-DNA conjugated antibodies allows efficient, multiplexed, spatialomic profiling of human and murine tissues

**DOI:** 10.1101/2020.11.16.385237

**Authors:** Anugraha Rajagopalan, Ishwarya Venkatesh, Rabail Aslam, David Kirchenbuechler, Shreyaa Khanna, David Cimbaluk, Vineet Gupta

**Affiliations:** Drug Discovery Center, Department of Internal Medicine, Rush University Medical Center, Chicago, Illinois, USA 60612; Department of Internal Medicine, Rush University Medical Center, Chicago, Illinois USA 60612; Center for Advanced Microscopy, Northwestern University Feinberg School of Medicine, Chicago, Illinois 60611; University of Illinois at Urbana-Champaign, Champaign, Illinois USA 61820; Department of Pathology, Rush University Medical Center, Chicago, Illinois USA 60612; Division of Hematology, Oncology and Cell Therapy, Department of Internal Medicine, Rush University Medical Center, Chicago, Illinois, USA 60612

## Abstract

Spatial organization of molecules and cells in complex tissue microenvironments provides essential cues during healthy growth and in disease. Novel techniques are needed for elucidation of their spatial relationships and architecture. Although a few multiplex immunofluorescence based techniques have been developed for visualization of the spatial relationships of various molecules in cells and tissues, there remains a significant need for newer methods that are rapid, easy to adapt and are gentle during the cyclic steps of fluorescence staining and de-staining. Here, we describe a novel, multiplex immunofluorescence imaging method, termed SeqStain, that uses fluorescent-DNA labelled antibodies for immunofluorescence staining of cells and tissues, and nuclease treatment for de-staining that allows selective enzymatic removal of the fluorescent signal. SeqStain can be used with primary antibodies, secondary antibodies and antibody fragments, such as Fabs, to efficiently analyse complex cells and tissues in multiple rounds of staining and de-staining. Additionally, incorporation of specific endonuclease restriction sites in antibody labels allows for selective removal of fluorescent signals, while retaining other signals that can serve as marks for subsequent analyses. The application of SeqStain on human kidney tissue provided spatialomic profile of the organization of >25 markers in the kidney, highlighting it as a versatile, easy to use and gentle new technique for spatialomic analyses of complex microenvironments.

## INTRODUCTION

Understanding the molecular and cellular composition of tissues and their relative organization in three-dimensional space of a complex tissue microenvironment (spatialomic organization) is essential for obtaining fundamental insights in tissue biology, interplay between various molecules and cells and for more accurately determining changes due to disease or treatments. This is especially true in the case of cancer, where insights into the complex tumour microenvironment helps guide therapeutic choices (1). Similar profiling of the cells infiltrating a transplanted tissue helps predict graft survival (2). Techniques to quantify multiple molecules in a single sample, such a transcriptomics, proteomics, flow cytometry and mass cytometry have revolutionized the field by providing deep characterization of tissue composition, yet they lack information about spatial organization of the various molecules and cells (3, 4). Immunohistochemical (IHC) and immunofluorescence (IF) based imaging methods provide information about spatial distribution of molecules and cells in tissues, yet were limited to detecting only 3-6 analytes at a time in the past (5–9). Thus, a great deal of recent effort has focused on increasing the multiplexing capabilities of such imaging-based techniques, and several methodologies for detecting multiple antigens in a single tissue section are now being developed (10–18).

Given that methods using brightfield IHC carries a high translation potential (7), techniques have been developed for detecting multiple antigens on a single tissue section by using cycles of IHC staining and de-staining using organic solvents (14, 19–21). Similarly, methodologies using IF based imaging have been developed for multiplex imaging of tissues (15–18). For example, recently described CycIF, a cyclic immunofluorescence method, achieves multiplex staining of tissue sections by combining cycles of immunofluorescent staining with de-staining using fluorescence bleaching (17, 22), whereas the 4i method achieves multiplex imaging by eluting antibodies after each round of staining (18). These methods, while providing high multiplex IF capabilities, use harsh protocols to remove the stain in each cycle, which has the potential to harm sensitive tissues.

DNA tagged antibodies are also used widely in imaging and offer the combined benefits of the specificity of antibodies and the versatility of DNA oligonucleotides (23, 24). Indeed, such reagents have redefined single-cell genomics by multiplexing cells from different samples (25, 26). DNA tagged antibodies have also recently been used very elegantly for multiplexed IF imaging. The CODEX method combines DNA tagged antibodies and in-situ primer extension with fluorescently tagged nucleotides to achieve high-level multiplexing (15). Although CODEX aids in deeper understanding of the tissue architecture, its complex experimental setup and design impedes wider accessibility of the technique. In contrast, Immuno-SABER uses cycles of in-situ DNA hybridization and removal (primer exchange reaction) for multiplexed IF imaging with DNA tagged antibodies (16). However, it also requires a complex design of DNA oligonucleotides and set up for the exchange of DNA strands and concatemers. Thus, a need exists for rapid, mild, and easy to use method for multiplexed immunofluorescence imaging.

Endonucleases are enzymes that selectively cleave oligonucleotides either non-specifically (such as DNase I) or in a sequence specific fashion (such as restriction endonucleases). These enzymes are routinely used in molecular and cell biology laboratories (27) to cleave oligonucleotide sequences rapidly and under mild conditions that do not harm the cells and tissues. Here, we describe a multiplex immunofluorescence imaging technique, termed SeqStain (for Sequential Staining), where we use fluorescent-DNA labelled antibodies to selectively stain antigens on cells and tissues, and use nucleases to rapidly cleave fluorescent signal off of the antibodies in the de-staining step. We show that the method is versatile, easy to use, gentle and highly effective in a multiplex environment and can be rapidly adapted for use on a variety of cells and tissues.

## MATERIALS AND METHODS

### Antibodies and oligonucleotides

Primary antibodies, secondary antibodies, and affinity purified Fc-specific Fab fragments, as described in **Table S1**, were obtained from commercial sources. Any preservative, such as sodium azide or glycerol, was removed from the commercially obtained antibodies using 50 kDa molecular weight cut-off Amicon ultra filtration columns (Millipore, # UFC505096) and buffer exchanged into phosphate-buffered saline (PBS) buffer pH 7.2. Synthetic oligonucleotides (oligos) were obtained from IDT (Coralville, Iowa). Oligonucleotide sequences are presented in **Table S2**.

### Cell culture

RAW264.7 cells (mouse), Lewis Lung Carcinoma (LLC) cells (mouse), K562 cells (human) and HeLa cells (human) were obtained from ATCC (Manassas, VA) and were cultured as per manufacturer’s recommendations. K562 CD11b/CD18 cells and mouse podocytes have been previously described (28, 29). All cell lines were maintained according to the published procedures at 37°C with 5% CO2.

### Preparation of fluorescent-DNA conjugated antibodies (SeqStain antibodies)

#### Preparation of linker oligos for conjugation to antibodies

Linker oligos were activated and conjugated to antibodies using published protocols (30). Briefly, linker oligos with a 5’-terminal amine modification were obtained from IDT (Coralville, Iowa) and activated using sulfo-SMCC (Sulfosuccinimidyl-4-(N-maleimidomethyl)cyclohexane-1-carboxylate) reagent (Thermo Fisher, #A39268) according to manufacturer’s instructions. This SMCC-modified oligo was cleaned up and desalted using the 7KDa molecular weight cut-off Zeba columns (Thermo Fisher, # 89882) and used in conjugation reaction with the reduced antibodies.

#### Conjugation of antibodies to the linker oligos

Antibodies were chemically conjugated to SMCC-modified linker oligos using published protocols (30). Briefly, antibodies were reduced using TCEP (Tris(2-carboxyethyl)phosphine hydrochloride) (Thermo Fisher, # 77720) according to manufacturer’s instructions, purified using 50 kDa molecular weight cut-off Amicon ultra filtration columns (Millipore, # UFC505096) and buffer exchanged into phosphate-buffered saline (PBS) buffer pH 7.2. Next, SMCC-modified oligo was added to the reaction mixture (at a 1:25 molar ratio), incubated at 4°C for 1 hour and purified using Amicon ultra filtration columns. Conjugation efficiency was determined using SDS-PAGE gels, which showed an average of 2-5 linker oligos chemically conjugated to an antibody during a typical labelling experiment. Subsequently, the conjugated antibody was washed, quantified by measuring the intensity of bands on an SDS-PAGE gel, and stored in PBS containing 0.5M NaCl.

#### Hybridization of oligos to create fluorescent-DNA labelled antibodies

The linker oligo conjugated antibody was mixed with fluorescent-DNA complex to prepare fluorescent-DNA labelled antibody. *First*, the fluorescent-DNA complex was prepared in a separate tube by mixing a docking oligo (**Figure S1A**) that contains a sequence complementary to the linker oligo and a set of sequence repeats for binding fluorescently labelled oligos. The fluorescent oligos were designed to contain a fluorophore at its 3’-end. The mixture was annealed by heating at 85°C for 5 minutes and slow cooling to generate the complex. All the oligos were obtained from IDT (Coralville, Iowa), with the fluorescent oligos containing a single fluorescent label at its 3’ end. *Next*, the complex was mixed with linker oligo conjugated antibody at 45°C and slow cooled to 37°C. The reaction mix was kept at 37°C for 30 minutes, slow cooled to room temperature and the completion of reaction was monitored by running it on a 4% agarose gel. Subsequently, the fluorescent-DNA labelled antibody complex was purified using 100 kDa molecular weight cut-off Amicon ultra filtration columns (Millipore #UFC510096) and stored in PBS buffer containing 0.5M NaCl at 4°C for use in SeqStain assays.

#### Preparation of SeqStain antibodies using DBCO-Azide chemistry

Antibodies were covalently conjugated to the linker oligo using the DBCO-Azide click chemistry using published protocols (31). Briefly, purified linker oligos containing 3’-terminal Azide were purchased from IDT (Coralville, Iowa) and resuspended in PBS, pH 7.2. Antibody was modified with DBCO-Sulfo-NHS ester (Click Chemistry Tools, #A124) according to manufacturer’s instructions. The DBCO activated antibodies were prepared and purified according to published protocols using 50 kDa molecular weight cut-off Amicon ultra filtration columns (Millipore, # UFC505096) and buffer exchanged into PBS, pH 7.2. DBCO-activated antibodies were combined with 20-molar fold excess of azide-modified linker oligo and incubated at 4°C overnight. Conjugation efficiency was estimated by running the mixture on a SDS-PAGE gel. The linker conjugated antibody was hybridized with fluorescent-DNA complex, as above, to prepare SeqStain antibodies.

#### Preparation of SeqStain antibodies using biotin-streptavidin methodology

Antibodies were biotin labelled in-house using a biotin labelling kit (Thermo Fisher, #A39257) by following manufacturer’s instructions. Linker oligo with 3’-terminal biotin was purchased from IDT (Coralville, Iowa) and purified streptavidin was purchased from BioLegend (Biolegend, #405150). Biotinylated antibody, streptavidin and biotinylated linker oligo were combined in a 1:1:3 molar ratio and incubated at room temperature for 30 minutes. Subsequently, the linker conjugated antibody was hybridized with fluorescent-DNA complex, as above, to prepare SeqStain antibodies.

### Preparation of fluorescent-DNA conjugated Fabs (SeqStain Fabs)

Affinity-purified Fc-specific Fab fragments were modified according to literature protocols using Traut’s reagent, which adds sulfhydryl groups by reacting with the primary amines in Fab (32, 33). Briefly, Traut’s reagent (Thermo Fisher, #26101) was added at 20-fold molar excess to the Fab fragments according to manufacturer’s instructions. This thiolation reaction proceeded for an hour at room temperature. Subsequently, excess reagent was quenched by adding 20mM glycine for 5 minutes. Thiolated Fab fragments were purified using 30 kDa molecular weight Amicon ultra filtration columns (Millipore, #UFC503096) and the concentration was estimated by A280 absorbance using Nanodrop 2000 (Thermo Fisher, Waltham, Massachusetts). Conjugation of the Fab with linker oligos and hybridization with fluorescent-DNA complex was performed as described for the antibodies above.

### Preparation of pre-complex between unlabelled primary antibody and SeqStain Fabs

SeqStain Fabs were mixed with unlabelled primary antibodies at equal molar ratio (3:1 weight ratio) in PBS, pH 7.2, and incubated at room temperature for 2 hours. Any unbound, excess Fab was removed by filtration using 100 kDa molecular weight cut-off size exclusion Amicon ultra filtration columns (Millipore, # UFC510096). Subsequently, the complex was used for staining of cells and tissue sections.

### Preparation of pre-complex between primary antibody and SeqStain secondary antibodies

Secondary antibodies were conjugated to the linker oligo using the maleimide-sulfhydryl chemistry and hybridized to the fluorescent oligo complex as detailed for primary antibodies. SeqStain secondary antibodies were mixed with unlabelled primary antibodies at 2:1 molar ratio at 0.1uM concentration with respect to the primary antibody to avoid formation of superclusters of primary and secondary antibodies. This reaction was incubated at room temperature for 15 minutes. Excess secondary antibody was then blocked by adding the corresponding normal serum at 1:20 v/v ratio and the mixture was incubated for 5 minutes at room temperature (Thermo Fisher Scientific, #10410 and #10510). The complex was used immediately for staining of cells and tissue sections.

### Gel analysis of SeqStain antibodies and Fabs

To determine linker oligo conjugation efficiency of antibodies, the reaction mixture was analysed using 10% SDS PAGE gel (Thermo Fisher Scientific, NW04122BOX) following manufacturer’s instructions. Controls, including unmodified antibodies and a protein ladder (Bio-rad, #1610375) were also used. Subsequently, the gel was Coomassie stained (Thermo Fisher Scientific, #24617) to visualize the protein bands. To determine efficiency of hybridization reaction, the annealing reaction mixture was analysed using a 4% agarose gel containing a DNA stain (Thermo Fisher Scientific, #S33102). The bands were visualized using a UV transilluminator (Bio-Rad laboratories, Hercules, California)

### Murine and human tissue samples

All animal studies were performed in compliance with the Institutional Animal Care and Use Committee (IACUC) at Rush University Medical Center. Murine spleen tissues were from mice bearing orthotopically transplanted LLC cells, as previously described (34). Briefly, LLC cells were passaged at least 3 times before inoculating into mice. Cells were routinely checked for mycoplasma using the MycoAlert assay (Lonza) and found negative prior to their use in animals. Wild type C57Bl/6 mice (6-8 week old) were obtained from The Jackson Laboratory. LLC cells (1.0 x 10^6^) in 100 μL cold phosphate-buffered saline (PBS) were subcutaneously inoculated into the mouse rear flank. Once the tumours grew to 1000mm^3^ in volume, the animals were sacrificed, and the harvested spleen tissue was immediately embedded in Tissue Tex OCT. Embedded tissue was cryopreserved in liquid nitrogen and transferred to −80°C for long term storage. Cryopreserved normal human kidney and tonsil tissue blocks were purchased from Origene (Rockville, MD).

### Flow cytometry assays

Single cell suspensions of cells in culture were prepared in PBS containing 1% BSA. Non-specific antibody recognition sites on cells were blocked by adding Fc Block (Biolegend, #101301) to the suspension for ten minutes at 4°C. Next, cells were stained with either unlabelled primary antibodies (conventional staining) or with SeqStain antibodies for 20 min at 4°C. Subsequently, cells were washed with PBS containing 1% BSA and analysed using LSR-Fortessa flow cytometer. Data was analysed using FlowJo software version 10.2.

### Staining of cells and tissue sections using SeqStain

#### Buffers and reagents

The following buffers and reagents were used in the assays. Block-1 solution (PBS containing 1% Bovine Serum Albumin (Sigma-Aldrich, #B4287-1G)), Block-2 solution (PBS containing 200ng/ml salmon sperm DNA (Thermo Fisher, #AM9680) and 3 nanomoles/ml singlestranded DNA and 0.5M NaCl), Wash buffer (PBS containing 0.1% Tween-20 (Sigma-Aldrich, #P1379), DNase I de-staining buffer (PBS containing 1X DNase buffer and 20 units of DNase I (NEB, #M0303) in 500ul), EcoRV de-staining buffer (deionized water containing 1X cut-smart buffer and 200 units of EcoRV (NEB, #R3195S) in 500ul), SmaI de-staining buffer (de-ionized water containing 1X cut smart buffer and 200 units of SmaI (NEB, #R01041S) in 500ul).

#### Preparation of cells for SeqStain

25 mm round cover glass (Fisher Brand, #12-545-102 25CIR-1) was coated with 3-Triethoxysilypropylamine APES (Millipore, #A3648) following manufacturer’s instructions. RAW264.7 cells were cultured as a monolayer on the coated cover glass in a 6-well plate. When the cells were about 50% confluent, they were removed from culture and fixed by treating with 4% paraformaldehyde at room temperature for 15 minutes. Once fixed, the cells were washed three times with PBS pH 7.4, permeabilized with 0.5% Triton-X at room temperature for 15 minutes. Subsequently, the cells were washed twice with PBS pH7.4 and blocked with the block-1 solution at room temperature for 1 hour. During this time, the cover glass was mounted onto the perfusion chamber.

#### Preparation of tissue sections for SeqStain

Tissue sections of 5μm thickness were mounted on the APES coated cover glass and fixed with acetone at room temperature for 5 minutes. Sections were further fixed by treating with 1.6% paraformaldehyde at room temperature for 15 minutes. The fixed sections were washed thrice with PBS pH7.4 and blocked with block-1 solution at room temperature for 1 hour. During incubation, the cover glass was mounted onto the perfusion chamber.

#### SeqStain immunofluorescence staining

The cover glass containing cells or tissue sections were blocked with Block-2 solution at room temperature for 1 hour. Subsequently, these samples were stained with either SeqStain antibodies or primary antibodies pre-complexed with SeqStain Fabs diluted in Block-2 solution at room temperature for 1 hour. Next, the chamber was perfused with the wash buffer continuously for 5 minutes to remove non-specific staining. Nuclei were counterstained by adding DAPI (Sigma-Aldrich, #D9542). The cells were washed again by perfusing wash buffer for 5 minutes and imaged. To de-stain, the chamber was perfused with 500ul of the de-staining buffer (NEB, #M0303) and imaged at the indicated time points. The de-staining buffer was replaced after 10 minutes for an additional round of de-staining. The de-stained samples were blocked again with block-2 solution for 20 minutes before adding the next round of SeqStain antibodies or primary antibodies pre-complexed with SeqStain Fabs. For the selective de-staining experiment, de-staining was performed by perfusing the appropriate de-staining buffers. Nuclear staining with DAPI was repeated every round. For staining by indirect immunofluorescence (conventional), the cover glass containing the cells or tissue sections were prepared as described before and mounted onto the perfusion chamber. The mounted samples were stained with primary antibodies after blocking with block-1 solution for 1 hour at room temperature. The chamber was perfused with wash buffer for 5 minutes to remove non-specific staining and then incubated with the corresponding secondary antibodies and stained at room temperature for 30 minutes. The samples were washed by perfusing wash buffer for 5 minutes and imaged after nuclear staining with DAPI.

#### Perfusion set-up

Cover glass was mounted on the closed bath chamber (Warner instruments, #RC-43C) following manufacturer’s instructions. This, in-turn, was mounted on the quick exchange platform (Warner instruments, #64-0375 (QE-1)) with built-in perfusion and suction holders. Buffers and solutions were perfused in and suctioned out via the inlet and outlet ports, respectively. A stage adapter (Warner instruments, #64-2415) was used to place the platform on the microscope to perform the iterative rounds of staining and de-staining.

#### Image acquisition

Immunofluorescence images were acquired using the Zeiss 700 LSM confocal microscope and Zen software (Carl Zeiss Group, Hartford, Connecticut). The destaining videos were acquired using the time-lapse image acquisition option in the Zen software. DNase I was added after the first image was acquired (−20 sec), following which images were acquired every 20 seconds for 5 minutes to monitor the rate of destaining. For the whole slide imaging, the images were acquired using the Nikon Eclipse TI2-E inverted confocal microscope (Nikon Corporation, Tokyo, Japan) with a motorized stage. The image tiles were acquired with the perfect focus system (PFS) to correct for any focal drifts between the tiles and then processed with the NIS Elements software.

#### Image analysis

Cell profiler version 3.1.9 was used to identify individual cells and their nuclei for both immunofluorescent staining of cells and tissue sections according to published protocols (35–37). Briefly, a working pipeline was created to run the required analysis. A set of modules were designed to identify objects and their relative intensities. For each image set, nuclei were considered as starting points and was identified using identify primary objects module. Identify secondary objects module was used to identify objects such as cells based on objects identified in previous module. Measure object intensity module was used to measure the intensity of each identified object. Overlay Object module created a color-coded label of the previously identified objects. Overlay Outline was used to place outlines of an object on a desired image as to create segmentation in the cellular region. Intensity measured was exported to excel files for further analysis.

For the whole slide images obtained from the multiplex imaging experiment, to align the images, DAPI images from each round of staining and de-staining were rigid registered (translation + rotation) to each other using the Register Virtual Stack Slices plugin in Fiji (38). For the other channels in each round, the same transformation as the corresponding DAPI channel was applied with the Transform Virtual Stack Slices plugin in Fiji. To measure distances of various cell types from the B-cell cluster, a region of interest was selected, and the cluster was traced manually using the signal from the DAPI and B220 channels. Euclidean Distance Map to the boarder of the cluster was calculated with Fiji. To measure signal intensities of each channel, the nuclei were first segmented using the DAPI channel. Segmentation was performed using the StarDist plugin, which uses an already trained model for convolution neural network (DSB 2018 dataset) (39). Once segmented, intensities for each channel was calculated on a per cell basis. Alongside, the EDM measurements for each segmented cell was calculated. In order to generate the plot profiles of the same markers from distinct rounds, first a stack of the corresponding images was generated. From the stack, a random region was cropped, and stack was converted to individual images. Plot profiles were calculated by drawing an arbitrary yellow line at the same position in both the images using Fiji. The overlay plots were generated using GraphPAD PRISM 9.

### Statistics

Statistical analyses were performed using Excel (Microsoft, Bothell, WA) and Prism 8.2 software (GraphPad Software). Student’s t-test were utilized when the data were normally distributed. Mann-Whitney test was used for comparison between two groups when data was not normally distributed.

## RESULTS

### A sequential combination of staining with fluorescent-DNA conjugated antibodies and destaining with nuclease provides a novel, easy-to-use, multiplexed imaging platform - SeqStain

DNA-tagged antibodies have recently been utilized in various techniques to perform multiplexed immunofluorescence imaging of cells and tissues (15–17, 22, 40, 41). These techniques rely on different methods for applying and removing fluorescent tags from the antibodies immobilized on substrates in an antigen specific fashion (15, 16, 42, 43). However, methods for cyclical removing or quenching of fluorescent labels and application of new ones is currently cumbersome and laborious, which also necessitates specialized equipment and setup. There is a current need for techniques that can perform rapid and selective cleavage of fluorescent labels from antibodies, that are also easy to use in a standard laboratory setting. Here, we describe an easy to use, rapid, multiplex imaging technique (termed “SeqStain”) that uses antibodies conjugated with fluorescently labelled DNA oligonucleotides (termed “SeqStain antibodies”). The method relies on sequential steps of immunofluorescent labelling with a set of such antibodies, gentle removal of the fluorescent labels post imaging, followed by another round of labelling with a new set of fluorescently labelled antibodies (**Figure 1A**). Importantly, the SeqStain methodology utilizes efficient and selective enzymatic processes, such as treatment with DNase I or restriction enzymes, for rapidly cleaving fluorescently labelled oligonucleotides (oligos) off of the antibodies, thereby removing the fluorescent labels from immune-fluorescently labelled substrates in the de-staining steps. Subsequently, the cleaved labels are washed off prior to initiation of the next round of staining. The enzymatic removal of fluorescent oligonucleotides also offers flexibility in the oligo sequence design, the length and complexity of the oligos and the types of oligos that can be used. Thus, cyclic steps of staining tissues with SeqStain antibodies and de-staining with nuclease treatment allows for efficient and rapid multiplex immunofluorescent imaging of cells and tissues.

**Figure 1.**
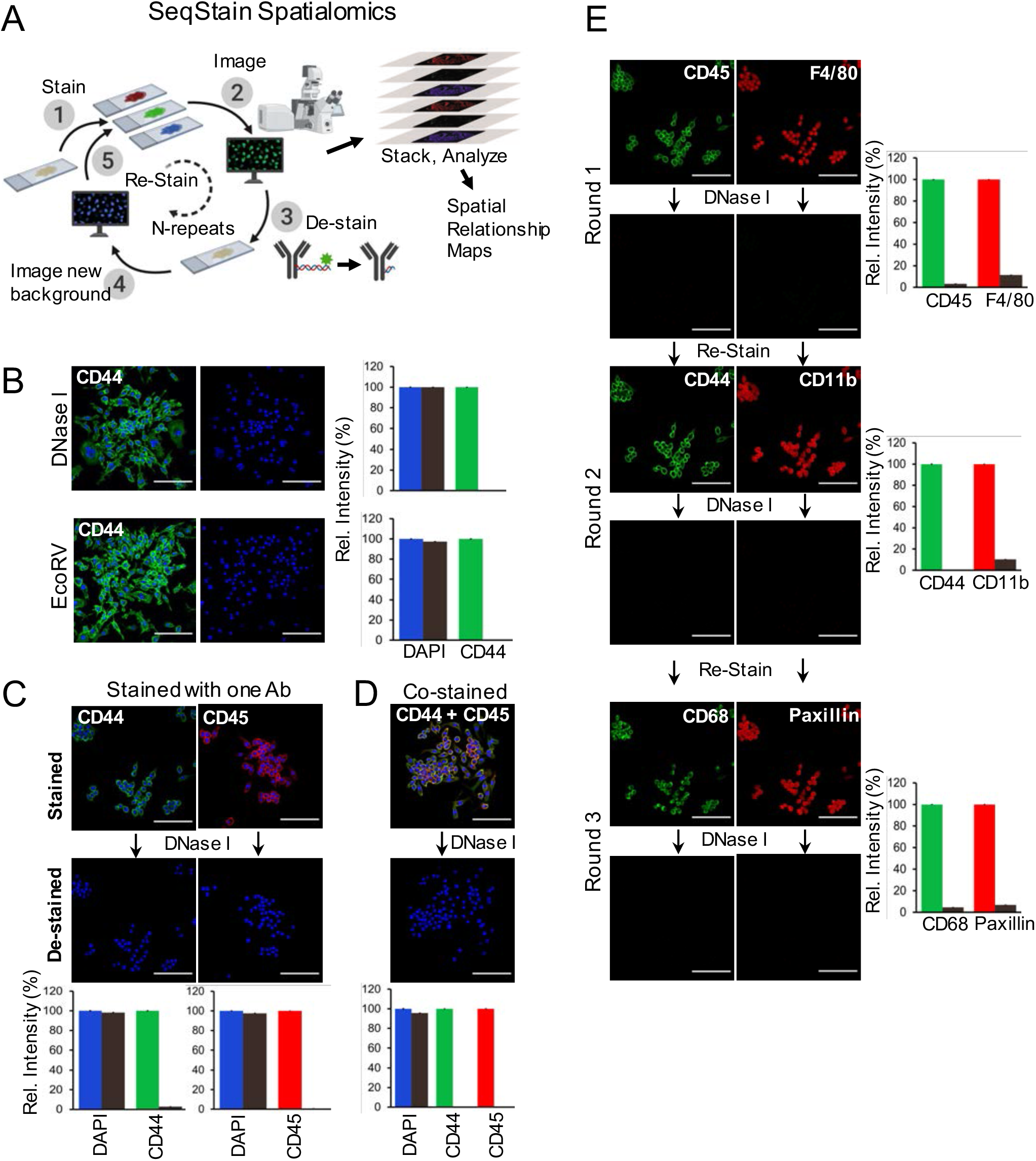
SeqStain based multiplex immunofluorescence imaging. **A**. Schematic representing the SeqStain methodology. Immobilized cells and tissue sections on glass are processed in cycles of immuno-staining with fluorescent-DNA labelled antibodies (step 1), imaging (step 2), gentle destaining using a nuclease (step 3), re-imaging (step 4) followed by next round of staining steps (step 5). Post-imaging, the data is analysed by computational stacking and alignment of the images followed by analyses of the spatial relationships between various markers to generate spatial maps of molecules and cells. Schematics were generated using Biorender. **B**. Immunofluorescence images of RAW264.7 cells after staining with anti-CD44 SeqStain antibody and after de-staining with either DNase I (top panels) or the endonuclease EcoRV (bottom panels). All images are representative of at least three replicates. Scale bar is 100μm. A bar graph showing quantification of fluorescence intensity for each panel is presented on the right. Graphs show the mean ± standard deviation. **C.** Immunofluorescence images of RAW264.7 cells after staining with anti-CD44 (fluorescently labelled with AF488 fluorophore) or anti-CD45 (labelled with Cy3 fluorophore) SeqStain antibodies (top panels) and after de-staining with DNase I for 1 min (bottom panels). Nuclei were labelled using DAPI. All images are representative of at least three replicates. Scale bar is 100 μm. A graph showing quantification of fluorescence intensity in each panel is presented on the bottom. Graphs show the mean ± standard deviation. **D.** Immunofluorescent images of RAW264.7 cells co-stained with anti-CD44 and anti-CD45 SeqStain antibodies (top panel) and 1 minute after the addition of DNase I (bottom panel). All images are representative of at least three replicates. Scale bar is 100μm. A graph showing quantification of fluorescence intensity in each panel is presented on the bottom. Graphs show the mean ± standard deviation. **E.** Immunofluorescence images of RAW264.7 cells after each of the three cycles of staining with two unique SeqStain antibodies and de-staining with DNase I. The antibodies used in each round are indicated in the panel, with SeqStain antibodies labelled using the AF488 fluorophore shown in green and the antibodies labelled using the Cy3 fluorophore shown in red. All images are representative of at least three replicates. Scale bar is 100μm. A graph showing quantification of fluorescence intensity after staining (green and red bars) and de-staining (brown bars) in each panel is also presented. Graphs show the mean ± standard deviation.

### SeqStain antibodies are highly efficient in multiplexed staining of mammalian cells

SeqStain antibodies were designed such that they could carry multiple fluorophores on each of the conjugated oligonucleotides (oligos) on the antibody (**Figure S1A**). Additionally, the oligo chains were designed so that the fluorescent dye molecules on the DNA were spaced apart by >5 nm dye-to-dye distance (at least 15 nucleotides apart) to prevent any unwanted dye-dye interactions (such as self-quenching) (44). SeqStain antibodies were generated in a two-step process (**Figure S1B**). In step 1, a 5’-amine modified 29-nucleotide long single-stranded DNA chain (linker oligo), containing a prespecified 15-mer sequence for hybridization with a complementary docking oligo, was covalently attached to the antibody using published protocols and purified (23, 31, 45, 46). Conjugation efficiency was determined using SDS-PAGE gels, which showed an average of 2-5 linker oligos chemically conjugated to an antibody during a typical labelling experiment (**Figure S1C**). In step 2, a fluorescently labelled double-stranded DNA (dsDNA) complex, containing a docking oligo and multiple fluorescently labelled oligos was hybridized to the antibody-DNA conjugate. This design of the dsDNA also allows for using many copies of fluorescently labelled DNA oligos to tune the signal intensity on the antibody. Subsequently, fluorescent-DNA labelled antibodies (SeqStain antibodies) were purified using the 100 kDa size-exclusion column. Formation of the fluorescent-DNA antibody complex was confirmed using 4% agarose gels (**Figure S1D**). This method of using fluorophores on the complementary oligonucleotides also avoids chemical quenching of the fluorophores during antibody-DNA conjugation step.

Flow cytometric analyses showed comparable staining of cell surface proteins CD45 and CD11b on RAW murine macrophage cells with SeqStain antibodies as compared to conventional staining using commercial antibodies (**Figure S2A**), confirming that antibody modification with DNA did not affect their ability to bind antigens. Additionally, staining of cells with the fluorescent oligo complex alone (no antibody) showed no staining. Similarly, immunofluorescent imaging of these cells with both SeqStain and conventional antibody labelling methods showed no difference between immunofluorescence labelling using DNA-modified primary antibody versus unmodified antibody (**Figure S2B**), and control staining with the fluorescent oligo complex alone showed no cellular staining. Furthermore, counterstaining of non-fluorescent SeqStain antibody labelled cells with a fluorescent secondary antibody showed cell staining similar to the staining observed with the same unmodified primary antibody stained using the conventional approach (**Figures S2C**), suggesting that the chemical conjugation of antibodies with oligonucleotides has minimal effect on their binding properties. Furthermore, oligos containing five repeats for complementary strand binding showed high fluorescence signal to background ratios (data not shown), and was, thus, utilized in all the assays presented here. Overall, this suggests that fluorescent dsDNA labelled primary antibodies provide signal intensity similar to the conventional approaches for use in immunofluorescence applications.

Next, to determine the feasibility of the technique, RAW cells were immobilized on glass cover slips and labelled with anti-CD44 SeqStain antibody. Of note, we incorporated an EcoRV restriction site in the dsDNA sequence on the SeqStain antibodies. After fluorescence imaging, treatment of cells with DNase I resulted in rapid loss of the fluorescence signal (**Figure 1B, top panels**). Additionally, treatment with EcoRV also resulted in rapid loss of fluorescence on the cells (**Figure 1B, bottom panels**), suggesting that both types of nucleases can be used in this methodology. Treatment of cells that were immunofluorescently stained with either a single SeqStain antibody (**Figure 1C**) or co-stained with two different SeqStain antibodies (**Figure 1D**) with DNase I rapidly reduced the fluorescence signal to background levels, suggesting that fluorescence signal from multiple fluorophores can be simultaneously removed in this technique. Furthermore, analysis of the timecourse for the de-staining step showed that treatment of stained samples with DNase I removed the fluorescence on sub-second timescales from both cells and human kidney tissue sections (**Supplementary Movies 1-4**). Keeping DNase I longer on the cells did not show any loss in sample integrity or increase in background signal (**Figure S3**). Furthermore, as chemical conjugation of linker oligo to antibody can be performed using a variety of methods, we tested two additional orthogonal chemistries to generate SeqStain antibodies (**Figures S4A-S4B)** (31, 47). We found that these different methods had no effect on the immunofluorescent staining and de-staining steps of SeqStain (**Figures S4C-S4D**), suggesting that modification of antibodies with fluorescent dsDNA can be successfully accomplished using multiple different methods.

Finally, to evaluate SeqStain technique for multiplex staining, a 6-plex panel with a combination of immune markers and structural markers was used to iteratively stain and de-stain the immobilized RAW cells in 3 rounds of staining and de-staining (**Figure 1E**). Staining was performed with a set of two unique antibodies, each with a unique fluorophore-labelled DNA tag, per round. Cell nuclei were stained using DAPI, to visualize nuclear boundary of each cells and to count cells. Fluorescent signal from the antibodies was removed using DNase I. Quantification showed complete removal of both fluorophores after each round, with no effect on DAPI staining, suggesting that this treatment does not affect the integrity of cells, nuclei or nuclear materials. A concern for the antibody based multiplex staining methods is that application of so many antibodies, either together or sequentially, could crowd the antigens, making multiplexing difficult. To address this concern in SeqStain, whether earlier round of staining may mask the antigens for probing at the later rounds, we stained and de-stained immobilized cells with the same SeqStain antibody for five cycles (**Figure S5**) and found no loss in immunofluorescent labelling capability of these samples or sample integrity. Moreover, the samples were further stained, de-stained and re-stained with a second set of two antibodies for five more rounds, without any loss of labelling capability or sample integrity. Together, these data show that SeqStain offers a rapid and robust multiplex immunofluorescence imaging platform.

### Primary antibodies pre-complexed with secondary antibodies and secondary antigen-binding fragments (Fabs) can also be used for multiplex staining by SeqStain

Conventional immunofluorescence imaging methods utilize antigen recognition using an unlabelled primary antibody followed by antibody recognition using a fluorescently labelled secondary antibody. However, extension of such methodology in a multiplex environment is significantly limited by the number of species (such as mouse, rat, goat, sheep) available to generate unique secondary antibodies. Pre-mixing primary antibodies with fluorescently labelled secondary antibodies or purified Fabs (the regions of antibodies with antigen binding capacity (48, 49)) prior to their use in staining could potentially overcome some of these limitations. Here, as an alternative to using primary SeqStain antibodies for multiplex staining, we tested if fluorescent-DNA labelled secondary antibodies (SeqStain secondary Ab) or Fabs of secondary antibodies (SeqStain Fabs) (**Figure 2A-2B**) could be applied in the SeqStain protocol. Such reagents have at least two major advantages. *One*, by precomplexing these reagents with primary antibodies, we can achieve signal amplification for targets with low level of expression. *Two*, these reagents may be quickly pre-complexed with many different primary antibodies, thus circumventing the need for modifying primary antibodies with fluorescent-DNA.

**Figure 2.**
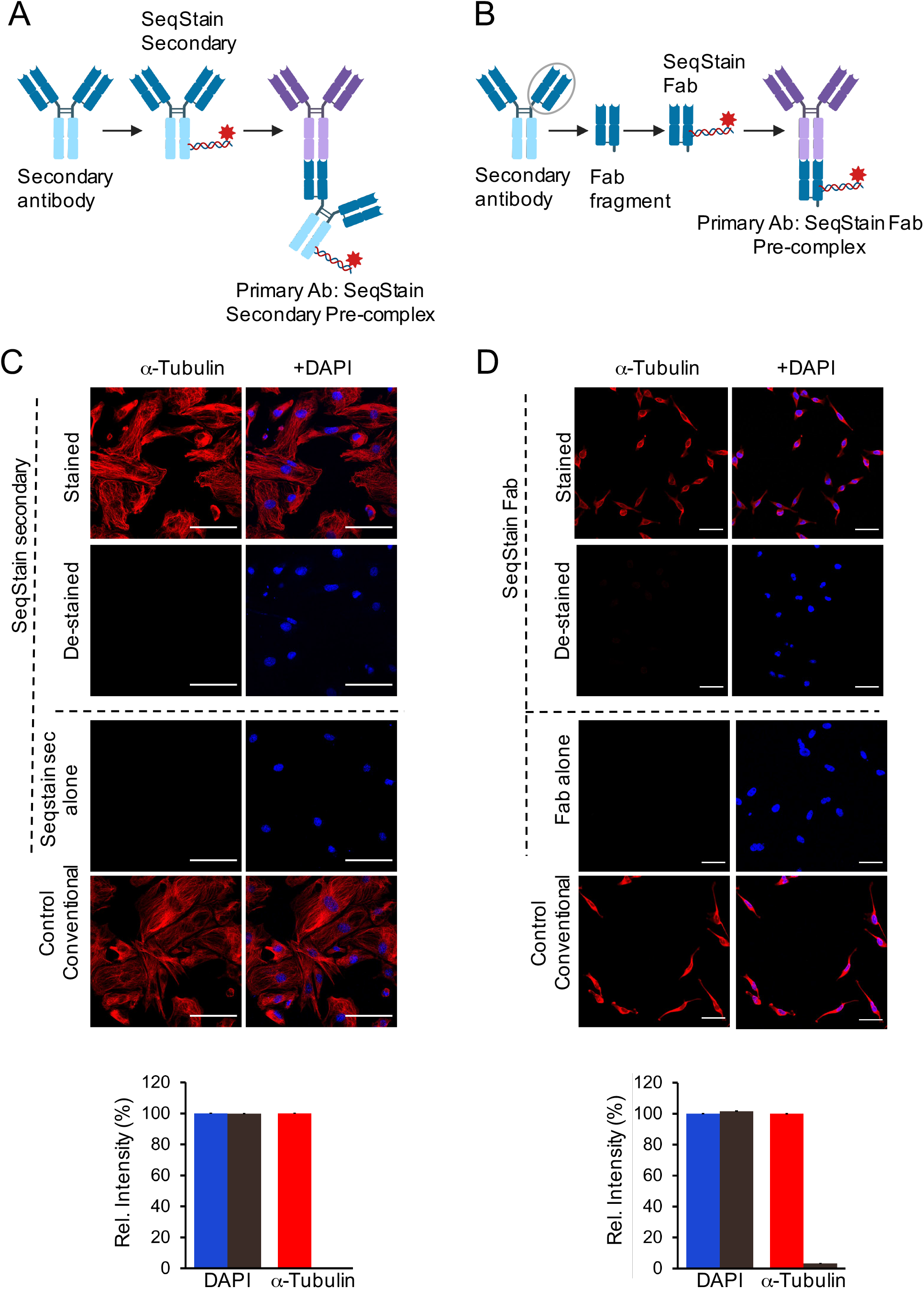
Application of SeqStain secondary antibodies and SeqStain Fabs in multiplex staining. A. Schematic representation of SeqStain secondary antibody based multiplex imaging method. Anti-Fc secondary antibodies are labelled with fluorescent DNA to develop SeqStain secondary antibodies. Subsequently, the SeqStain secondary antibodies are pre-complexed with appropriate primary antibodies and used in staining. **B.** Schematic representation of SeqStain Fab based multiplex imaging method. Fab fragments generated by enzymatic digestion of antibodies are labelled with fluorescent DNA to develop SeqStain Fabs. Subsequently, SeqStain Fabs are pre-complexed with their corresponding primary antibodies for use in multiplex staining. **C**. Immunofluorescence images of murine podocyte cell line stained with anti a-tubulin antibody precomplexed with SeqStain secondary antibody (left panels) and with DAPI nuclear staining (right panels). Immunofluorescent images after destaining with DNase I are shown below each panel. Control staining of these cells with SeqStain secondary alone and with conventional methodology is shown in the bottom panels. All images are representative of at least two replicates. Scale bar is 100μm. A graph showing quantification of fluorescence intensity in each panel is also presented. Graphs show the mean ± standard deviation. **D**. Immunofluorescence images of HeLa cells stained with anti a-tubulin antibody pre-complexed with SeqStain Fab (left panels) and with DAPI nuclear staining (right panels). Immunofluorescence images of these cells after destaining with DNase I are shown below each panel. Control staining of these cells with SeqStain Fab alone (secondary Fab) and with conventional methodology is shown in bottom panels. All images are representative of at least two replicates. Scale bar is 100μm. A graph showing quantification of fluorescence intensity in each panel is also presented. Graphs show the mean ± standard deviation.

To test, we prepared SeqStain Fabs using commercially available secondary antibodies or with affinity purified Fc-specific Fab fragments from secondary antibodies (mouse, rat and rabbit) (**Figure S6**). Primary rabbit anti-mouse antibody against a-Tubulin was pre-complexed with anti-rabbit SeqStain secondary Ab or with anti-rabbit secondary SeqStain Fab, respectively. Next, we stained immobilized mouse podocytes (**Figure 2C**) or HeLa cells (**Figure 2D**) with them. Immunofluorescence imaging showed high level of staining similar to the staining with conventional primary-secondary antibodies. Expectedly, treatment of these SeqStain secondary Ab or the SeqStain Fab stained cells with nuclease DNase I resulted in rapid loss of the fluorescence signal. As a control, staining with SeqStain Secondary Ab or SeqStain Fab alone, in the absence of the primary rabbit anti-mouse antibody against a-Tubulin did not show any staining, highlighting the specificity of this approach. Furthermore, use of a 6-plex panel of primary antibodies pre-complexed with SeqStain Fabs to iteratively stain and de-stain immobilized RAW cells also showed efficient labelling and clearing of fluorescent signal from these cells (**Figure S7**), further suggesting that SeqStain methodology can be applied with a variety of reagents for rapid multiplex immunofluorescence imaging.

### SeqStain allows rapid, multiplexed immunofluorescence imaging of complex tissues

To test the feasibility of the SeqStain approach on complex tissues, we tested it on murine (spleen) and human tissues (kidney, tonsil) (details in **Methods and Materials** section). We selected a set of commercially available, well-characterized antibodies (**Table S1**) that recognize several different cell types for these assays. *First*, we validated the prepared SeqStain antibodies for appropriate pattern of staining using conventional protocols and literature provided results. Data in **Figure 3** show that each of the SeqStain reagent provided expected antigen staining patterns on tissues comparable to the conventional immunofluorescence staining assays. For example, staining of mouse spleen sections (**Figure 3A**) showed that mouse anti-IgD and anti-IgM SeqStain antibodies expectedly stained B cell clusters and the anti-CD169 SeqStain antibody stained marginal zone macrophages around the B cell cluster in ring-like pattern (15, 50, 15) similar to the staining observed using conventional methods. Staining using anti-CD68 and anti-CD11b SeqStain antibodies, that stain macrophages in the spleen, provided expected staining patterns comparable to that of the conventional method. Remarkably, we also found that staining of serial sections of the same murine spleen tissue for an antigen (MHC II) using two different approaches - SeqStain and conventional immune-staining, respectively - yielded similar pattern of MHC II positive B cell clusters throughout the tissue section (52) (**Figure 3B**), suggesting that SeqStain antibodies provide results similar to conventional labelling approaches with primary and secondary antibodies. Similarly, staining of human kidney tissue sections (**Figure 3C**) showed that anti-human SeqStain antibodies against Cytokeratin-8, CD31, CD45, Synaptopodin, Podocin, Na+K+-ATPase, EpCAM and Collagen IV provided expected and comparable staining patterns as the unmodified antibodies used in conventional approach. Not surprisingly, use of SeqStain Fab pre-complexed with primary antibodies also provided staining pattern results comparable to the staining patterns observed using conventional indirect immunofluorescence, as exemplified on human kidney and tonsil tissue sections in **Figures 3D and 3E**. Furthermore, we found that the type of fluorophore used to label the dsDNA complex on SeqStain antibodies did not have any material effect on their performance, suggesting that most commonly available fluorophores can be used to fluorescently tag the DNA on the antibodies (**Figure S**8). Together, these data suggest that SeqStain antibodies and Fabs are highly efficient in immunostaining of a variety of complex tissues and provide staining results like those obtained with conventional immunofluorescence staining methods.

**Figure 3.**
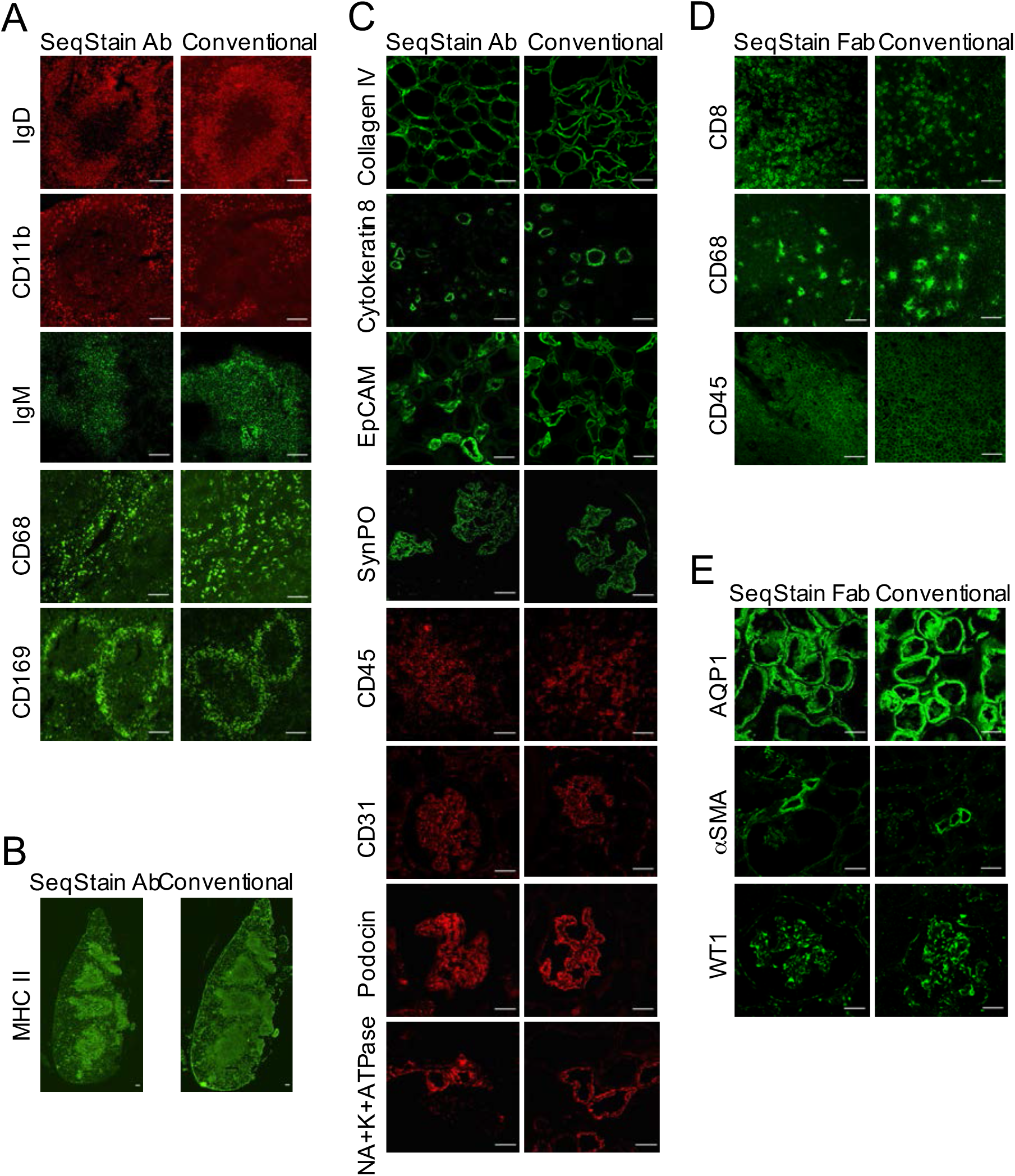
Comparison of SeqStain based staining with conventional immunofluorescence staining of various tissues. **A.** Immunofluorescence images showing mouse spleen tissue stained using SeqStain antibody based method (SeqStain Ab) or using conventional immunostaining method (conventional). The antibodies used are indicated in the panel, with antibodies labelled using the AF488 fluorophore shown in green and the antibodies labelled using the AF546 fluorophore shown in red. All images are representative of at least three replicates. Scale bar is 100μm. **B.** Immunofluorescence images showing whole slide scan of serial sections of mouse spleen tissue stained for MHC II using either the SeqStain antibody method (left panel) or the conventional method (right panel). Scale bar is 100μm. **C**. Immunofluorescence images showing human kidney tissue stained using SeqStain antibody based method (SeqStain Ab) or using conventional immunostaining method (conventional). The antibodies used are indicated in the panel, with antibodies labelled using the AF488 fluorophore shown in green and the antibodies labelled using the AF546 fluorophore and Cy3 shown in red. All images are representative of at least three replicates. Scale bar is 100μm. **D.** Immunofluorescence images showing mouse spleen tissue stained using SeqStain Fab based method (SeqStain Fab) pre-complexed with a primary antibody or using conventional immunostaining method (conventional). The antibodies used are indicated in the panel and were stained using the AF488 fluorophore. All images are representative of at least three replicates. Scale bar is 100μm. **E.** Immunofluorescence images showing human kidney tissue stained using SeqStain Fab based method (SeqStain Fab) pre-complexed with a primary antibody or using conventional immunostaining method (conventional). The antibodies used are indicated in the panel and were stained using the AF488 fluorophore. All images are representative of at least two replicates. Scale bar is 100μm.

Subsequently, we evaluated the feasibility of using enzymatic approach for removing the DNA-linked fluorophore in tissues immunostained with SeqStain antibodies. Surprisingly, we found that nucleases were equally efficient in removing the fluorescent signal from SeqStain antibody stained tissue sections as they were with immobilized cells, even though the tissues provide a highly complex microenvironment. Human kidney tissue (, **Figure 4**) stained with a variety of SeqStain antibodies showed a complete removal of fluorescence signal after a brief treatment with DNase I. Notably, DNase I treatment of tissue sections stained using the fluorescent antibodies in the conventional method (where the fluorophores do not contain a DNA linker when they are chemically linked to the antibodies) did not result in any loss of signal (**Figure S9**). More importantly, DNase I treatment did not harm the integrity of the tissues. To probe this further, especially to make sure that the extensive DNase I treatment will not remove DNA binding proteins from the nuclear DNA, we treated human kidney sections with increasing amounts of DNase I and subsequently stained the cells with anti-Histone H1 antibody. Results show no loss in Histone levels or chromosomal DNA, suggesting that these treatments do not reduce sample quality (**Figure S10**).

**Figure 4.**
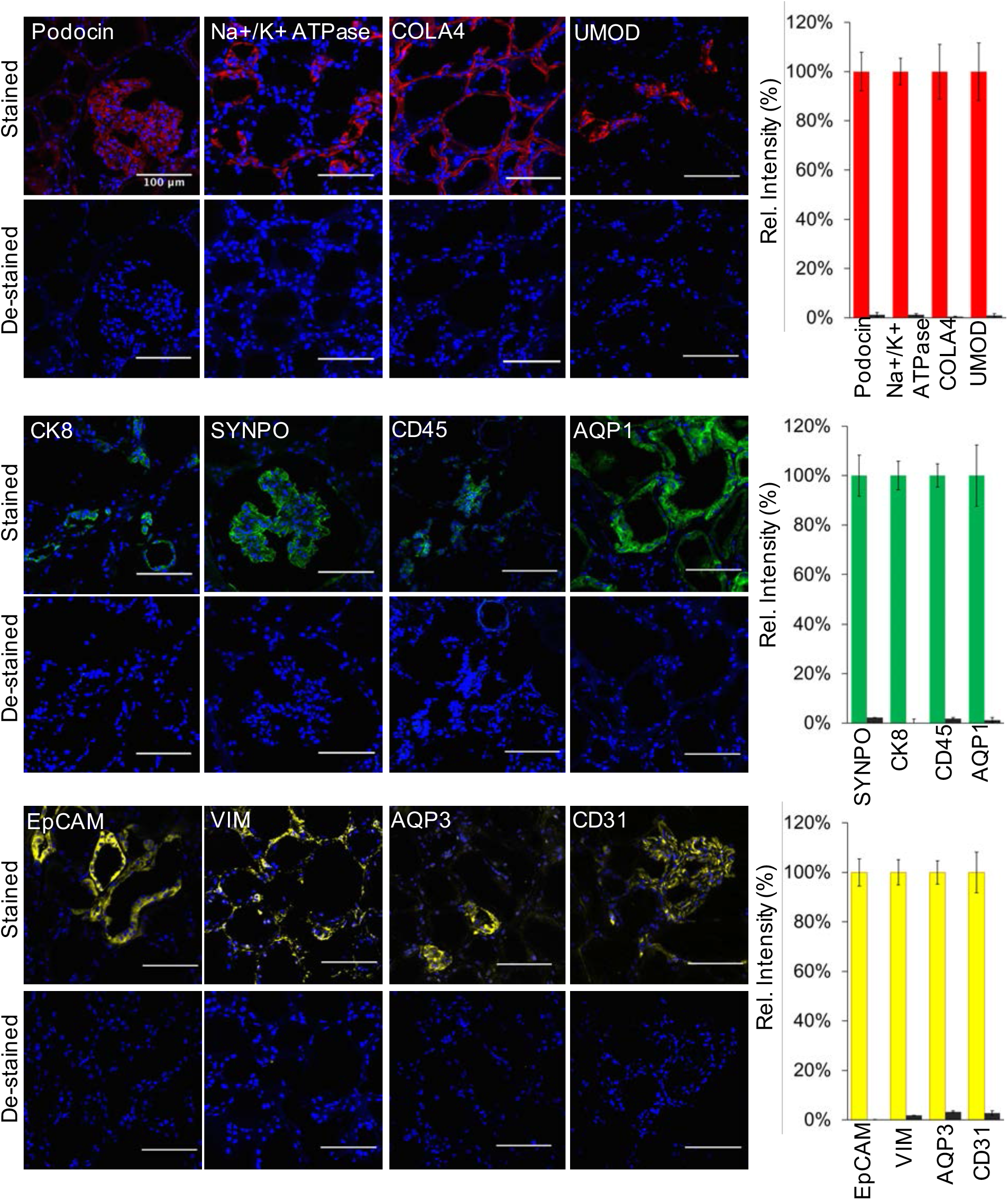
Enzymatic de-staining of tissues stained using the SeqStain antibodies. Immunofluorescence images showing human kidney tissue sections stained using SeqStain antibodies (as indicated in the panel). The antibodies were labelled using either the AF488 fluorophore (shown in green), the Cy3 fluorophore (shown in red) or the Cy5 fluorophore (shown in yellow). Immunofluorescence images of these tissue sections after de-staining with DNase I treatment are shown below each panel. All images are representative of at least three replicates. Scale bar is 100μm. Graphs showing quantification of fluorescence intensity after staining (red bars, green bars or yellow bars) and de-staining (brown bars) in each panel is also presented on the right. Graphs show the mean ± standard deviation.

Finally, when tissue sections stained with SeqStain antibodies were co-stained with conventional fluorescent secondary antibody, using spectrally non-overlapping fluorophores, it showed overlapping staining pattern (**Figure S11**), suggesting that the chemical modification of primary antibody with fluorescent dsDNA sequences does not harm the antibody properties and does not hamper its recognition by a secondary antibody. Again, treatment of the doubly labelled tissue section with DNase I showed selective removal of the SeqStain fluorophore, without negatively affecting the conventionally conjugated fluorophore, the staining pattern, or the tissue integrity. But when the SeqStain secondary antibodies pre-complexed with primary antibodies were used, DNase I treatment completely removed the fluorophore signal from the secondary antibodies. (**Figure S12**), Together, these data comprehensively show that SeqStain methodology is equally efficient for immunofluorescence staining of complex tissues, provides providing a novel method for gentle and rapid removal of the fluorescent signal from tissues and has high promise for use in multiplex tissue imaging.

### Fluorescent labels from tissues labelled using SeqStain antibodies can be selectively removed using pre-determined choice of nucleases

It is often desirable to maintain immunofluorescent label on an antigen, while staining and de-staining other antigens on the same population of cells and tissues. However, such applications are difficult to incorporate using the current multiplex imaging techniques. Given that SeqStain relies on nuclease-based enzymatic removal of fluorescent signals, SeqStain methodology offers the possibility of selective removal of fluorophores, while maintaining fluorescent label of another fluorophore during cycles of multiplex imaging. To study the feasibility of this approach, we engineered two unique restriction sites (EcoRV and SmaI) in the linker oligo sequences and used them to prepare SeqStain antibodies (**Figure 5A**). Thus, we prepared SeqStain antibodies against EpCAM and Collagen IV that were labelled with fluorescent DNA containing EcoRV restriction site and SeqStain antibodies against Na+K+-ATPase and Aquaporin1 that were labelled with fluorescent DNA containing SmaI restriction site. Subsequently, we labelled human kidney cryosections with a combination of anti-Na+K+-ATPase and anti-EpCAM SeqStain antibodies bearing AF488 and Cy3 fluorochrome tag, respectively (**Figure 5B**). Imaging results showed expected labelling of the distinct tubular segments with the two different antibodies(15, 53). Next, to selectively remove only one of the two fluorescent labels, we treated the tissue samples with EcoRV restriction endonuclease, that resulted in selective removal of the signal from the anti-EpCAM, but did not affect the fluorescent signal from the anti-NA+K+-ATPase antibody (**Figure 5B**). Subsequently, to determine if this would have any negative consequences on staining with antibody labelled with DNA containing the same EcoRV restriction site and to study the utility of this method in a multiplexed setting, in the next round of staining, we applied anti-Collagen IV SeqStain antibody containing EcoRV restriction site and the Cy3 fluorescent tag. Imaging results show tubular basement membrane stained by the Collagen IV antibody along with the Na+K+-ATPase positive transport channels in the tubules that were labelled in the previous round of staining (**Figure 5B**). Next, for selective cleavage of the other signal, we treated the tissue sample with SmaI restriction endonuclease, which selectively removed the signal from anti-Na+K+-ATPase antibody. Finally, to determine if this would have any negative consequences on staining with antibody labelled with DNA containing an SmaI site, in the next round, we applied anti-Aquaporin1 antibody containing SmaI restriction site and the AF488 fluorophore, with expected staining of the cortical proximal tubules upon imaging (**Figure 5B**). The results clearly show that the SeqStain methodology allows selective de-staining of fluorescent markers in a multiplex setting offering significant advantages where maintaining staining of specific markers or antigens is needed. Such retained stains can also serve as structural markers for imaging in subsequent rounds and to help better orient the stacks of the whole tissue during image analyses.

**Figure 5.**
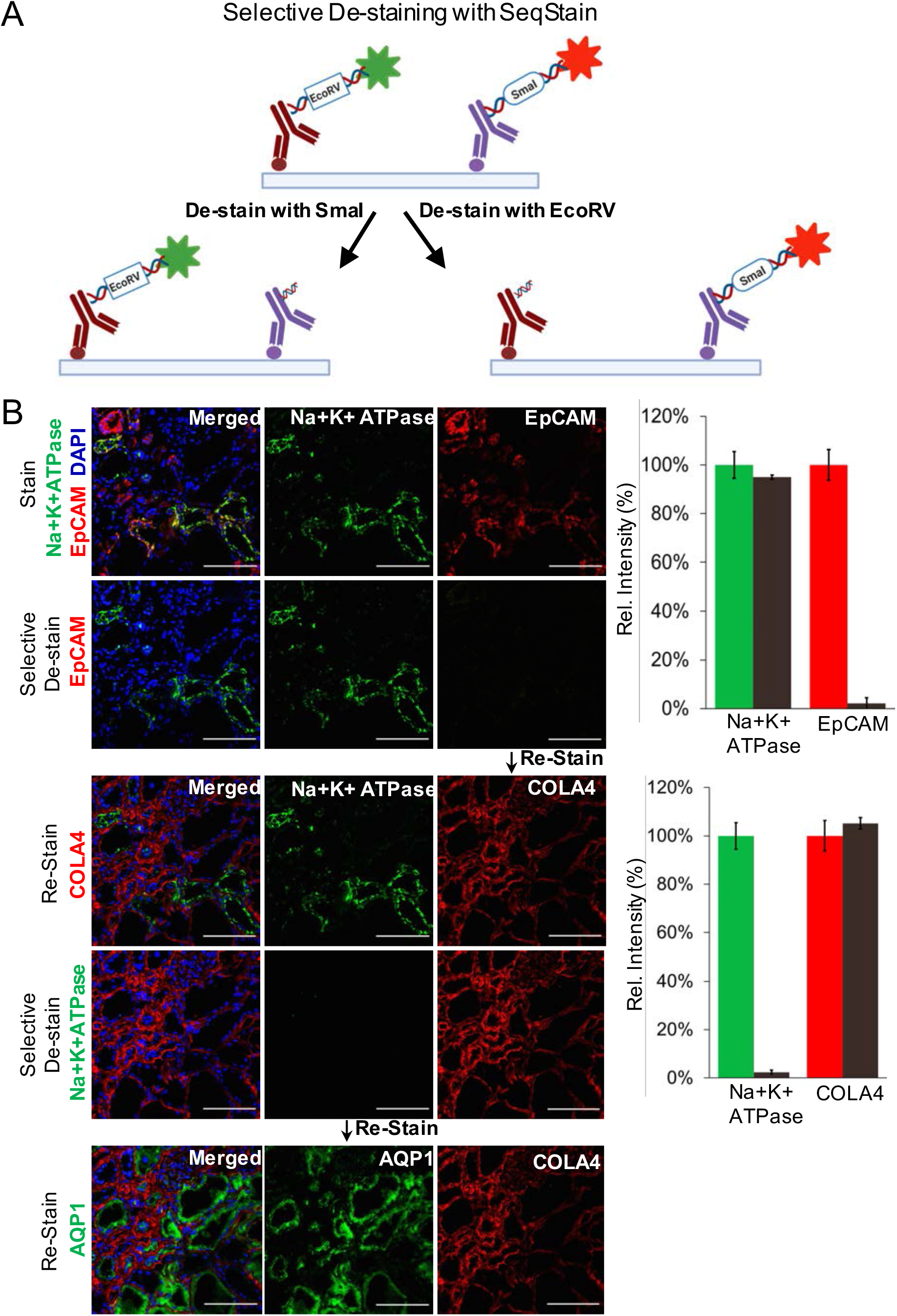
Selective de-staining of complex tissues using SeqStain antibodies and restriction endonucleases. **A.** Schematic representation of the technique for selective removal of fluorophores from immunofluorescently labelled tissues is shown. Tissue can be stained with a combination of SeqStain antibodies labelled using fluorescent DNA that contains specific recognition sites for restriction endonuclease(s) (such as EcoRV and SmaI). Treatment of samples with the specific restriction nuclease selectively removes fluorophores only from the antibodies carrying the respective DNA sequence, leaving all others undiminished. **B.** Immunofluorescence images showing normal human kidney tissue sections after each of the cycles of staining with unique SeqStain antibodies (as indicated in the panel) and de-staining with a specific restriction endonuclease (as indicated). The antibodies were labelled using the AF488 fluorophore (shown in green) or the Cy3 fluorophore (shown in red). Merged images from the two fluorescence channels (along with images from DAPI stained nuclear markers) are also shown. All images are representative of at least three replicates. Scale bar is 100μm. Graph showing quantification of fluorescence intensity after staining (green and red bars) and de-staining (brown bars) in each panel is also presented on the right. Graphs show the mean ± standard deviation.

### SeqStain enables spatialomic profiling of multiple antigens on spleen and kidney tissues

To test spatialomic profiling via multiplex immunofluorescence staining of a single tissue section using SeqStain, in a proof-of-concept experiment, we developed a panel of nine unique antibodies against various immune markers, along with pan-nuclear marker DAPI, and used it in five rounds of sequential staining and de-staining steps on mouse spleen tissue. Each round of staining used two unique SeqStain antibodies followed by imaging of the whole slide and de-staining with nuclease

DNase I. Again, DAPI-stained nuclei provided a guidepost for aligning the various image sets at the end of the experiment. Results in **Figure 6A** show high level of fluorescence staining by each SeqStain antibody, followed by complete and rapid removal of fluorescent label from both channels after DNase I treatment (**Figure S13**). After completion of the imaging rounds, images were stacked and aligned using DAPI stained nuclei using ImageJ (38, 54). Cell Profiler based image analyses showed that the individual cells can be efficiently segmented computationally for cell-based fluorescent signal analyses of SeqStain stained tissues (**Figure S14**).

**Figure 6.**
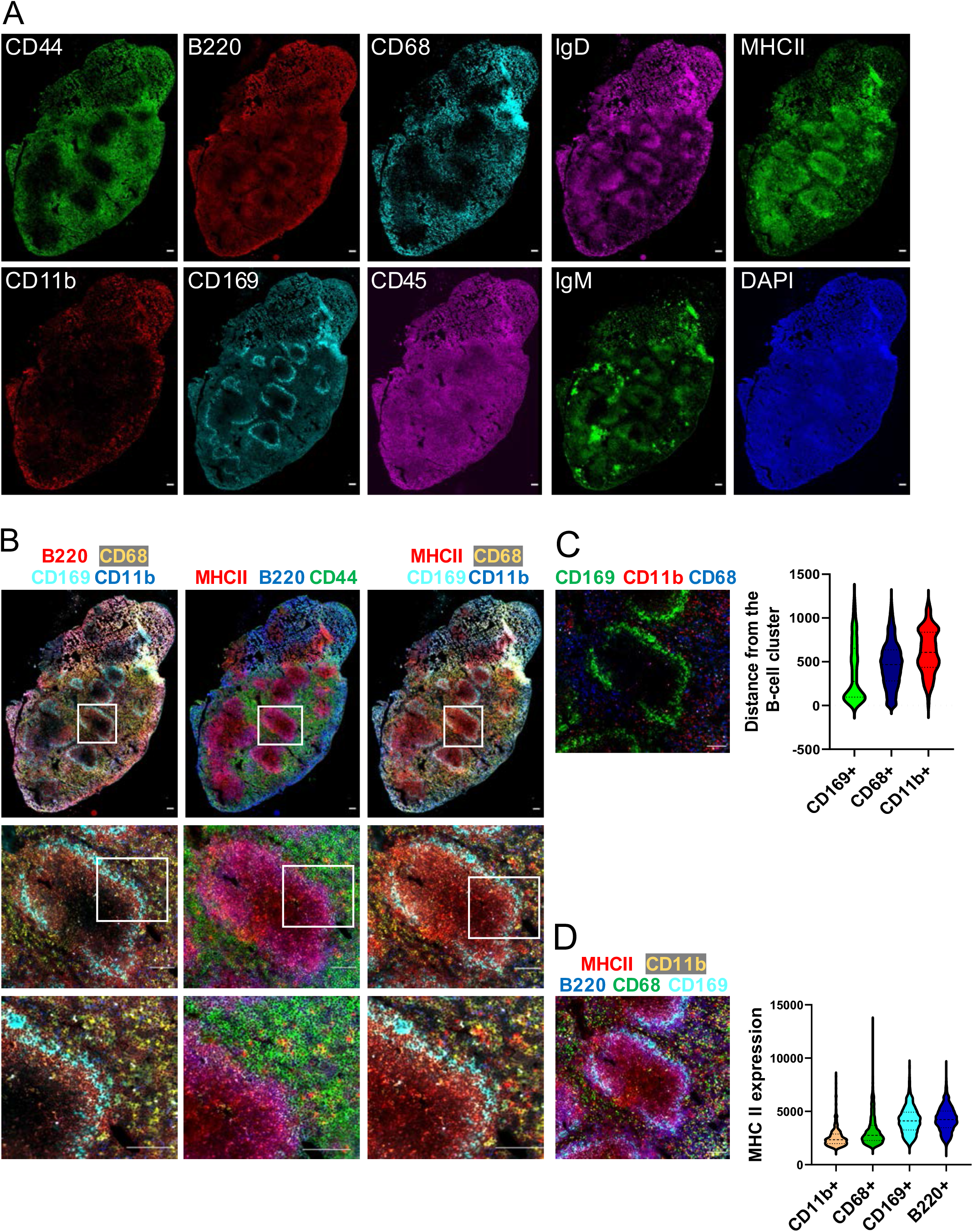
Multiplex imaging of whole spleen tissue using a 9-plex SeqStain panel. **A.** Immunofluorescence images of whole mouse spleen tissue sections after multiplex staining with multiple rounds of staining with unique SeqStain antibodies and DAPI (as indicated in the panel). Scale bar is 100μm. **B-D.** Composite overlays of aligned immunofluorescence image stacks from whole mouse spleen tissue sections after multiplex staining with SeqStain antibodies and DAPI (as indicated on top of each panel). (Below) zoomed-in regions from each composite immunofluorescence image, showing co-localization of various markers based on SeqStain staining. Scale bar is 100μm **C.** Composite overlay showing the location of various myeloid cells (as indicated on the top of the panel) with respect to a selected B-cell cluster. Scale bar is 100μm. Violin plot on the right shows the computed distance for each myeloid cell type on a per cell basis from the B-cell cluster. Graphs show the mean ± standard deviation. **D.** Composite overlay showing MHC II expression on various cell types (as indicated on the top of the panel) in a selected region. Scale bar is 100μm. Violin plot on the right shows computed co-expression of MHC II and the indicated markers on a per cell basis. Graphs show the mean ± standard deviation.

Subsequently, select composites were generated from the aligned images to show the spatial organization of different markers in the spleen tissue (**Figure 6B-6D**). Images of the entire tissue section clearly showed no changes in overall tissue morphology or integrity during due to the repeated cycles of staining and de-staining. Staining of different myeloid cells using CD169, CD68 and CD11b revealed their spatial relationship with respect to B220+ B cells. Expectedly, the CD169+ macrophages were found in the marginal zone area lining B220+ B cell clusters, in close contact with the B-cells as is typical for these cell population whereas the CD68+ macrophages were predominantly found in the red pulp region of the spleen. Additionally, the CD11 b+ cells were found scattered outside the red pulp region (53, 55, 56). Thus SeqStain can be used to profile the heterogenous myeloid populations in the spleen whose distinct spatial location and relationship with other cell types help orchestrate the immune response during an infection (57, 58).

We also characterized the MHC II expression and on various cell types in the spleen tissue and their respective spatial relationship. Quantification of MHC II levels revealed that CD68+ and CD11b+ macrophages in the spleen have low levels of MHC II expression (59). While CD169+ macrophages residing in the marginal zone area had higher expression of MHC II (60, 61). Expectedly, B220+ B cells had the highest expression of MHC II among the cell types analysed (62, 63). Thus, by quantifying the relative spatial location of cells and their co-expression of markers, SeqStain offers a simple yet robust method to understand the spatial organization of various cell types in the tissue and an ability to generate spatial relationship maps (SRMs).

Next, we expanded antibody set to profile human kidney tissue and also tested the feasibility of using three-color antibody mixtures for staining in each round of SeqStain. We developed a 20-color panel (19 unique antibodies + DAPI) and used it to stain a human kidney tissue section in 9 cycles of staining and de-staining (**Figures 7 and S15**). This time we used three unique SeqStain antibodies in each round of staining, followed by imaging of the whole tissue section, de-staining with nuclease DNase I and re-imaging. Additionally, to confirm staining, we used a few of the antibodies twice, to obtain a 25-plex image of the kidney tissue at high resolution (**Figure S15**). As above, images were stacked and aligned using DAPI stained nuclei in ImageJ for data quantification (38, 54). Results show high level of immunofluorescence staining by each SeqStain antibody, followed by complete and rapid removal of fluorescent label from each of the three channels after DNase I treatment. Aligned images of the whole tissue section clearly showed no loss of integrity of the tissue of changes in tissue morphology due to the repeated cycles of staining and de-staining. This was further confirmed from the analyses of images after repeated staining with the same antibody in a different cycle, such as Vimentin and Aquaporin1. Analyses of the stained images from two different cycles clearly showed high overlap of fluorescence signal without any loss in signal intensity or increase in background fluorescence (**Figure S16**). Further analyses of various kidney tissue substructures using 7-plex images from this experiment clearly showed presence of expected cellular neighbourhoods in the various parts of the kidney (**Figure 8**). Analyses of various substructures in the renal cortex revealed their spatial relationships. Composites of pseudocolored images revealed a predominant presence of proximal tubules (AQP1) consistent with the histology of renal cortex (**Figure 8A**) (64, 65). We were also able to identify the collecting duct (AQP2+ AQP3+ EpCAM+) (Cyto7+ Cyto8+) (**Figure 8A and 8C**) (66–68), Distal convoluted tubules (AQP2-AQP3-EpCAM+), Thick Ascending Loop of Henle’s (EpCAM+ Uromodulin+) and their spatial location with respect to one another (**Figure 8A**) (69). Similarly, in the glomerulus, we were able to discern the three components of the glomerular capillary filter-podocytes, glomerular basement membrane and glomerular endothelial cells (69). Podocytes can be identified by the co-expression of WT1 and Vimentin, the glomerular basement membrane can be visualized by Collagen IV staining and the glomerular endothelial cells by staining for CD31 (**Figure 8B**). In addition, the specialized contractile mesenchymal cells of the glomerulus can be discerned by Vimentin staining in the absence of WT1 staining. In the renal interstitium (70–73), Collagen IV delineates the tubular basement membrane forming a network throughout the entire tissue (74). Peritubular capillaries visualized by CD31 staining can be seen as a disconnected network of cells within this tubular basement membrane (75). The renal interstitium also contains stromal cells which can be visualized by Vimentin and a-SMA staining (**Figure 8B**) (76). Angiotensin Converting Enzyme (ACE2), the functional receptor for the SARS coronaviruses, was found to be highly expressed in the proximal tubular cells identified by ACE2 coexpression with AQP1 (77, 78). The normal human kidney had resident immune cells which were identified by CD45 staining while the renal macrophages were identified by co-expression of CD45 and CD68 (**Figure 8C**) (79). In summary, these experiments show that SeqStain is highly applicable for multiplexed spatial profiling of various types of tissues and allows for rapid generation of spatial relationship maps.

**Figure 7.**
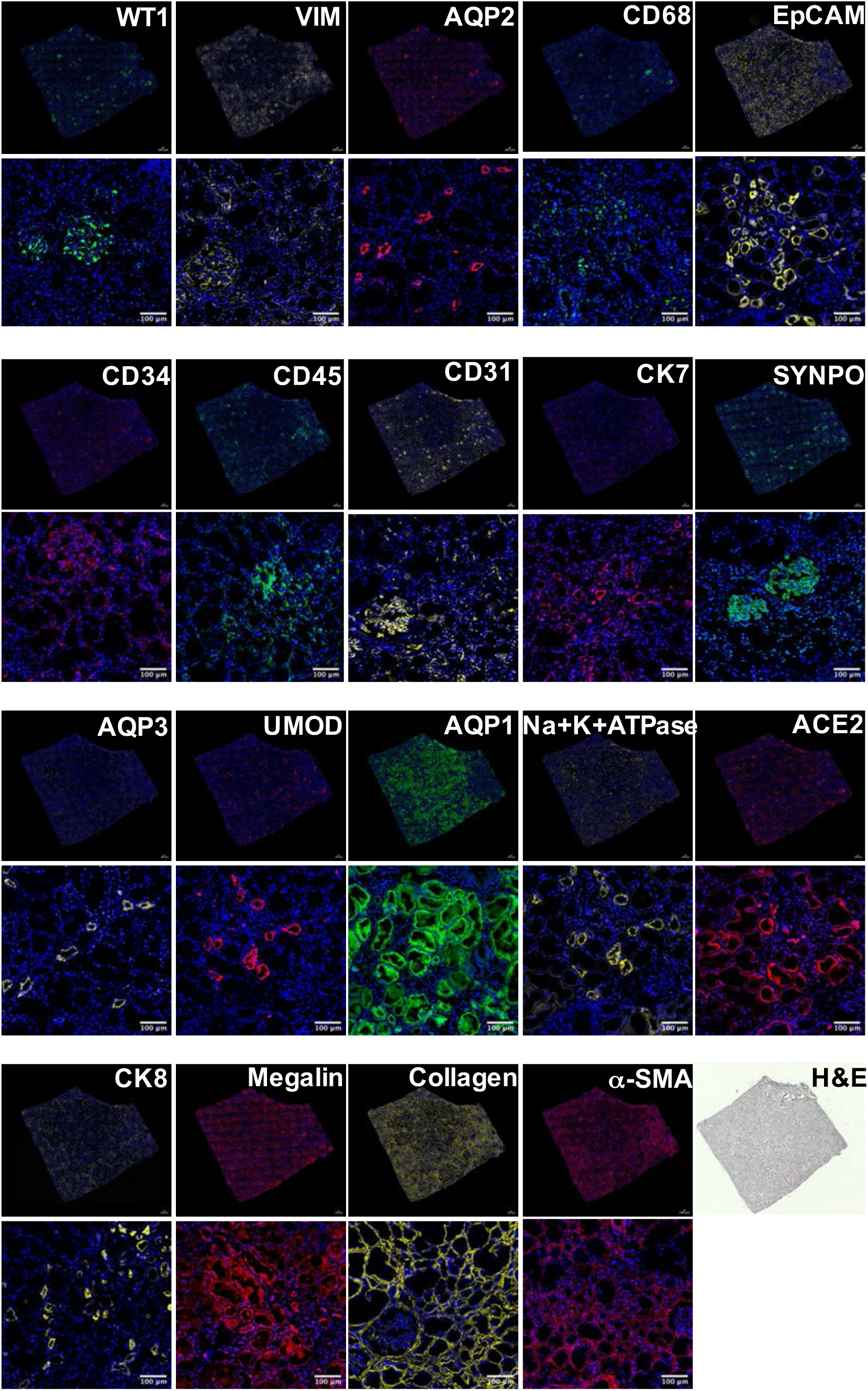
SeqStain based multiplex imaging of whole human kidney tissue provides a 20-plex image. Immunofluorescent images of whole kidney tissue sections after each round of staining with unique SeqStain antibodies (as indicated) and DAPI (as indicated in the panel). Zoomed-in sections of images are presented below each panel. Scale bar is 100μm. A serial section stained with H & E is also presented.

**Figure 8.**
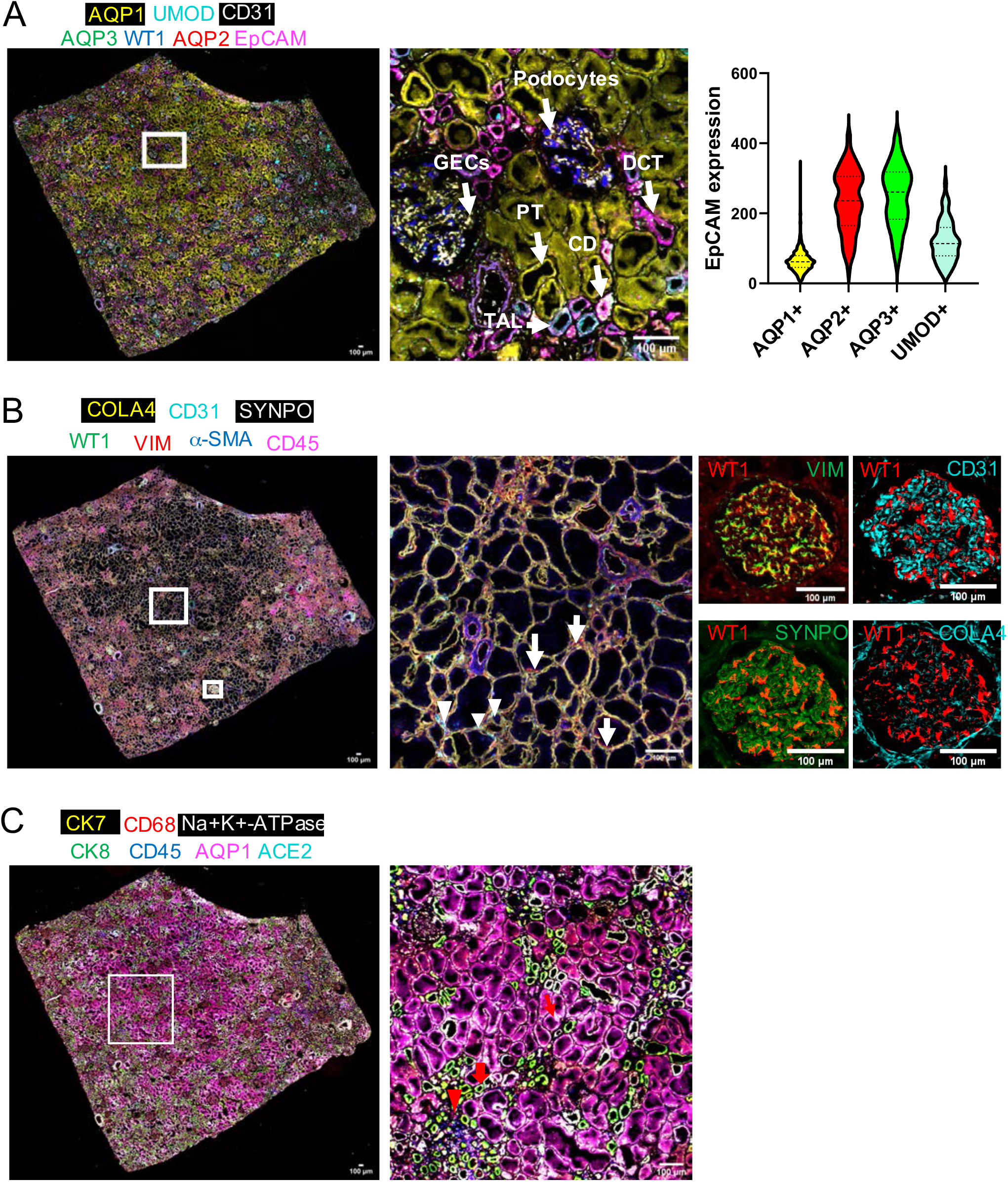
Multiplex imaged panels identify major substructures in the human kidney. **A**. Image showing a composite overlay of aligned immunofluorescence image stack of seven unique markers from the whole kidney tissue sections (left panel, markers as indicated on the panel). Image of the zoomed-in region in the middle panel shows various components of the kidney tissue, including the Glomerular Endothelial Cells (GECs), Proximal Tubule (PT), Collecting duct (CD), Distal Convoluted Tubules (DCT) and Podocytes. Violin plot on the right shows the computed co-expression of EpCAM and the indicated markers on a per cell basis. Graphs show the mean ± standard deviation. Scale bar is 100μm. **B**. Image showing a composite overlay of aligned immunofluorescence image stack of seven unique markers from the whole kidney tissue sections (left panel). Image of the zoomed-in region in the middle panel shows the renal interstitium (middle panel), long arrow points to fibroblasts and short arrow points to cortical peritubular capillaries. Images of a zoomed-in region show a renal glomerulus (four panels on the right), false coloured for the indicated markers. **C**. Image showing a composite overlay of aligned immunofluorescence image stack of seven unique markers from the whole kidney tissue sections (left panel). Image of the zoomed-in region (right panel) shows specific cell types and parts, including the renal macrophages (red arrow heads), collecting duct (thick red arrow), and the receptor ACE2 in proximal tubules (thin red arrow). Scale bar is 100μm.

These data also suggest that, although the experiments here utilized either a 10-plex or a 25-plex SeqStain panel to evaluate multiplexed staining, the method is easily scalable to tens of different markers. Signal amplification, if necessary for markers expressed at low levels, can be done using a number of published techniques (80–82), that include either increasing the number of fluorophores on the oligos, increasing the number of fluorescent oligos on the docking oligos, using different DNA structures (such as branched DNA (83), origami folded plasmids (43), Fluorocubes (84) etc) or by precomplexing primary antibodies with SeqStain Fabs and SeqStain secondary antibodies.

## DISCUSSION

Spatial profiling of cells in tissues provides critical insights into disease pathogenesis and can be diagnostic. The importance of such insights is appreciated especially for diseases like cancer where spatial heterogeneity often leads to poor clinical outcomes (1, 85, 86). Indeed, to address such needs, the recent years have seen a surge of innovative multiplex staining techniques at both transcriptomic and proteomic levels (10–19, 87, 88). However, the existing protein multiplexing methods either use harsh de-staining conditions or require complex experimental set-up. The SeqStain spatialomic analysis methodology presented here is a gentle, easy-to-use, and efficient multiplex imaging technique that provides a unique platform for obtaining such spatialomic insights. The method applies fluorescent DNA-labelled antibodies with gentle, enzymatic method for removing fluorescence signal after each cycle of staining. In particular, we demonstrate multiplex staining of immobilized cells and tissue sections, where de-staining gently removes the fluorescent signal to pre-staining levels on the whole slide. Strikingly, de-staining using the SeqStain method was also rapid. We observed that treatment with nucleases removed >99% of the signal in <1 minute, without affecting sample integrity or tissue morphology. Additionally, by engineering specific restriction sites into DNA during antibody modification, we show that selective de-staining is possible with SeqStain. Retention of selective markers for subsequent rounds may be important for spatial aligning of tissues when it is not possible to perform whole slide scanning or for measuring information about multiple neighbouring antigens via FRET or other such methods, where keeping fluorophore fixed on one antigen might be helpful, while changing the second or third fluorophores on different antigens. SeqStain is thus a highly configurable, multiplex staining method which uses the familiar laboratory essentials such as the endonucleases system to achieve rapid multiplexing of cells and tissues. Furthermore, the enzymatic approach for the removal of fluorescent labels offers flexibility in the design of oligo sequence used for conjugation, the length and complexity of the oligos, the types of fluorophores that can be included, and the types of oligo-based higher order structures that can be used.

The technique offers significant new advantages for deeper understanding of complex tissue microenvironments. This novel methodology uses commercially available primary and secondary antibodies and Fab fragments that are chemically conjugated with fluorescently labelled doublestranded DNA (SeqStain antibodies and Fabs) that are easy to modify in any laboratory. We also show that such modifications do not affect their function in any way by testing in a variety of systems. To avoid cross-reactivity, pre-complexation of SeqStain Fabs or SeqStain secondary antibodies with primary antibodies can also be used during staining. This paves way to build a multiplex panel that combines SeqStain antibodies, SeqStain secondary antibodies and SeqStain Fabs by opening up additional ways of staining tissues in multiplex analyses. Utilizing Fabs and secondary antibodies for SeqStain not only improves sensitivity of detection for markers expressed at low levels, but it also facilitates higher throughput by reducing the time it takes to prepare reagents for SeqStain. Furthermore, SeqStain is performed using a simple perfusion set-up with readily available components which together forms an elegant staining platform. In fact, this set-up transforms standard confocal microscopes into a multiplex imaging gear which when combined with a motorized stage enables high level data acquisition by this method.

In summary, spatialomic profiling of cells and tissues is crucial to discern the underlying molecular architecture and their various interactions. The SeqStain methodology presented here, uses cyclic steps of staining tissues with fluorescent DNA-labelled antibodies and de-staining with nuclease treatment, is a gentle, efficient, and rapid technique for multiplex immunofluorescent imaging of cells and tissues that has wide applicability.

## SUPPLEMENTARY DATA

Supplementary Data containing Figures S1-S16 and Movies 1-4 are available at NAR online.

## ACKNOWLEDGEMENT

We thank Liudmila Romanova (Mila) and Diptaman Chatterjee (Neil) in the Kordower laboratory at Rush University Medical Center, University of Illinois at Chicago (UIC) Histology Core and the Center for Advanced Microscopy at Northwestern University Medical Center for help with tissue staining and image analysis; Thomas Cunningham, Mohd. Hafeez Faridi and other past and current members of the Gupta laboratory for their invaluable technical help and suggestions during this project.

## REFERENCES

1. Binnewies, M., Roberts, E.W., Kersten, K., Chan, V., Fearon, D.F., Merad, M., Coussens, L.M., Gabrilovich, D.I., Ostrand-Rosenberg, S., Hedrick, C.C., et al. (2018) Understanding the tumor immune microenvironment (TIME) for effective therapy. Nat. Med., 10.1038/s41591-018-0014-x.

2. Sablik, K.A., Jordanova, E.S., Pocorni, N., Clahsen-van Groningen, M.C. and Betjes, M.G.H. (2020) Immune Cell Infiltrate in Chronic-Active Antibody-Mediated Rejection. Front. Immunol., 10.3389/fimmu.2019.03106.

3. Bodenmiller, B., Zunder, E.R., Finck, R., Chen, T.J., Savig, E.S., Bruggner, R. V., Simonds, E.F., Bendall, S.C., Sachs, K., Krutzik, P.O., et al. (2012) Multiplexed mass cytometry profiling of cellular states perturbed by small-molecule regulators. Nat. Biotechnol., 10.1038/nbt.2317.

4. Bendall, S.C., Nolan, G.P., Roederer, M. and Chattopadhyay, P.K. (2012) A deep profiler’s guide to cytometry. Trends Immunol., 10.1016/j.it.2012.02.010.

5. Gannot, G., Tangrea, M.A., Erickson, H.S., Pinto, P.A., Hewitt, S.M., Chuaqui, R.F., Gillespie, J.W. and Emmert-Buck, M.R. (2007) Layered peptide array for multiplex immunohistochemistry. J. Mol. Diagnostics, 10.2353/jmoldx.2007.060143.

6. Frampton, J.P., Tsuei, M., White, J.B., Abraham, A.T. and Takayama, S. (2015) Aqueous two-phase system-mediated antibody micropatterning enables multiplexed immunostaining of cell monolayers and tissues. Biotechnol. J., 10.1002/biot.201400271.

7. Peng, C.W., Liu, X.L., Chen, C., Liu, X., Yang, X.Q., Pang, D.W., Zhu, X.B. and Li, Y. (2011) Patterns of cancer invasion revealed by QDs-based quantitative multiplexed imaging of tumor microenvironment. Biomaterials, 10.1016/j.biomaterials.2010.12.053.

8. Tóth, Z.E. and Mezey, É. (2007) Simultaneous visualization of multiple antigens with tyramide signal amplification using antibodies from the same species. J. Histochem. Cytochem., 10.1369/jhc.6A7134.2007.

9. Cappi, G., Dupouy, D.G., Comino, M.A. and Ciftlik, A.T. (2019) Ultra-fast and automated immunohistofluorescent multistaining using a microfluidic tissue processor. Sci. Rep., 10.1038/s41598-019-41119-y.

10. Stack, E.C., Wang, C., Roman, K.A. and Hoyt, C.C. (2014) Multiplexed immunohistochemistry, imaging, and quantitation: A review, with an assessment of Tyramide signal amplification, multispectral imaging and multiplex analysis. Methods, 10.1016/j.ymeth.2014.08.016.

11. Dixon, A.R., Bathany, C., Tsuei, M., White, J., Barald, K.F. and Takayama, S. (2015) Recent developments in multiplexing techniques for immunohistochemistry. Expert Rev. Mol. Diagn., 10.1586/14737159.2015.1069182.

12. Fountaine, T.J., Wincovitch, S.M., Geho, D.H., Garfield, S.H. and Pittaluga, S. (2006) Multispectral imaging of clinically relevant cellular targets in tonsil and lymphoid tissue using semiconductor quantum dots. Mod. Pathol., 10.1038/modpathol.3800628.

13. Gerdes, M.J., Sevinsky, C.J., Sood, A., Adak, S., Bello, M.O., Bordwell, A., Can, A., Corwin, A., Dinn, S., Filkins, R.J., et al. (2013) Highly multiplexed single-cell analysis of formalinfixed, paraffin-embedded cancer tissue. Proc. Natl. Acad. Sci. U. S. A., 10.1073/pnas.1300136110.

14. Remark, R., Merghoub, T., Grabe, N., Litjens, G., Damotte, D., Wolchok, J.D., Merad, M. and Gnjatic, S. (2016) In-depth tissue profiling using multiplexed immunohistochemical consecutive staining on single slide. Sci. Immunol., 10.1126/sciimmunol.aaf6925.

15. Goltsev, Y., Samusik, N., Kennedy-Darling, J., Bhate, S., Hale, M., Vazquez, G., Black, S. and Nolan, G.P. (2018) Deep Profiling of Mouse Splenic Architecture with CODEX Multiplexed Imaging. Cell, 10.1016/j.cell.2018.07.010.

16. Saka, S.K., Wang, Y., Kishi, J.Y., Zhu, A., Zeng, Y., Xie, W., Kirli, K., Yapp, C., Cicconet, M., Beliveau, B.J., et al. (2019) Immuno-SABER enables highly multiplexed and amplified protein imaging in tissues. Nat. Biotechnol., 10.1038/s41587-019-0207-y.

17. Lin, J.-R., Fallahi-Sichani, M. and Sorger, P.K. (2015) Highly multiplexed imaging of single cells using a high-throughput cyclic immunofluorescence method. Nat. Commun., 6, 8390.

18. Gut, G., Herrmann, M.D. and Pelkmans, L. (2018) Multiplexed protein maps link subcellular organization to cellular states. Science (80-.)., 10.1126/science.aar7042.

19. Akturk, G., Sweeney, R., Remark, R., Merad, M. and Gnjatic, S. (2020) Multiplexed Immunohistochemical Consecutive Staining on Single Slide (MICSSS): Multiplexed Chromogenic IHC Assay for High-Dimensional Tissue Analysis. In Methods in Molecular Biology.

20. Huang, W., Hennrick, K. and Drew, S. (2013) A colorful future of quantitative pathology: Validation of Vectra technology using chromogenic multiplexed immunohistochemistry and prostate tissue microarrays. Hum. Pathol., 10.1016/j.humpath.2012.05.009.

21. Glass, G., Papin, J.A. and Mandell, J.W. (2009) SIMPLE: A sequential immunoperoxidase labeling and erasing method. J. Histochem. Cytochem., 10.1369/jhc.2009.953612.

22. Lin, J.-R., Izar, B., Wang, S., Yapp, C., Mei, S., Shah, P.M., Santagata, S. and Sorger, P.K. (2018) Highly multiplexed immunofluorescence imaging of human tissues and tumors using t-CyCIF and conventional optical microscopes. Elife, 7.

23. Kazane, S.A., Sok, D., Cho, E.H., Uson, M.L., Kuhn, P., Schultz, P.G. and Smider, V. V. (2012) Site-specific DNA-antibody conjugates for specific and sensitive immuno-PCR. Proc. Natl. Acad. Sci. U. S. A., 10.1073/pnas.1120682109.

24. Sano, T., Smith, C.L. and Cantor, C.R. (1992) Immuno-PCR: Very sensitive antigen detection by means of specific antibody-DNA conjugates. Science (80-.)., 10.1126/science.1439758.

25. Stoeckius, M., Zheng, S., Houck-Loomis, B., Hao, S., Yeung, B.Z., Mauck, W.M., Smibert, P. and Satija, R. (2018) Cell Hashing with barcoded antibodies enables multiplexing and doublet detection for single cell genomics. Genome Biol., 10.1186/s13059-018-1603-1.

26. Gaublomme, J.T., Li, B., McCabe, C., Knecht, A., Yang, Y., Drokhlyansky, E., Van Wittenberghe, N., Waldman, J., Dionne, D., Nguyen, L., et al. (2019) Nuclei multiplexing with barcoded antibodies for single-nucleus genomics. Nat. Commun., 10.1038/s41467-019-10756-2.

27. Roberts, R.J. (2005) How restriction enzymes became the workhorses of molecular biology. Proc. Natl. Acad. Sci. U. S. A., 10.1073/pnas.0500923102.

28. Maiguel, D., Faridi, M.H., Wei, C., Kuwano, Y., Balla, K.M., Hernandez, D., Barth, C.J., Lugo, G., Donnelly, M., Nayer, A., et al. (2011) Small molecule-mediated activation of the integrin CD11b/CD18 reduces inflammatory disease. Sci. Signal., 4.

29. Shankland, S.J., Pippin, J.W., Reiser, J. and Mundel, P. (2007) Podocytes in culture: Past, present, and future. Kidney Int., 10.1038/sj.ki.5002291.

30. Williams, B.A.R. and Chaput, J.C. (2010) Synthesis of peptide-oligonucleotide conjugates using a heterobifunctional crosslinker. Curr. Protoc. Nucleic Acid Chem., 10.1002/0471142700.nc0441s42.

31. Gong, H., Holcomb, I., Ooi, A., Wang, X., Majonis, D., Unger, M.A. and Ramakrishnan, R. (2016) Simple Method to Prepare Oligonucleotide-Conjugated Antibodies and Its Application in Multiplex Protein Detection in Single Cells. Bioconjug. Chem., 10.1021/acs.bioconjchem.5b00613.

32. Jue, R., Lambert, J.M., Leland, P.L., Pierce, R. and Traut, R.R. (1978) Addition of Sulfhydryl Groups to Escherichia coli Ribosomes by Protein Modification with 2-Iminothiolane (Methyl 4-Mercaptobutyrimidate). Biochemistry, 10.1021/bi00618a013.

33. Newton, D.L., Hansen, H.J., Mikulski, S.M., Goldenberg, D.M. and Rybak, S.M. (2001) Potent and specific antitumor effects of an anti-CD22-targeted cytotoxic ribonuclease: Potential for the treatment of non-Hodgkin lymphoma. Blood, 10.1182/blood.V97.2.528.

34. Schmid, M.C., Khan, S.Q., Kaneda, M.M., Pathria, P., Shepard, R., Louis, T.L., Anand, S., Woo, G., Leem, C., Faridi, M.H., et al. (2018) Integrin CD11b activation drives anti-tumor innate immunity. Nat. Commun., 9.

35. Carpenter, A.E., Jones, T.R., Lamprecht, M.R., Clarke, C., Kang, I.H., Friman, O., Guertin, D.A., Chang, J.H., Lindquist, R.A., Moffat, J., et al. (2006) CellProfiler: Image analysis software for identifying and quantifying cell phenotypes. Genome Biol., 10.1186/gb-2006-7-10-r100.

36. Lamprecht, M.R., Sabatini, D.M. and Carpenter, A.E. (2007) CellProfiler™: Free, versatile software for automated biological image analysis. Biotechniques, 10.2144/000112257.

37. McQuin, C., Goodman, A., Chernyshev, V., Kamentsky, L., Cimini, B.A., Karhohs, K.W., Doan, M., Ding, L., Rafelski, S.M., Thirstrup, D., et al. (2018) CellProfiler 3.0: Next-generation image processing for biology. PLoS Biol., 10.1371/journal.pbio.2005970.

38. Arganda-Carreras, I., Sorzano, C.O.S., Marabini, R., Carazo, J.M., Ortiz-De-Solorzano, C. and Kybic, J. (2006) Consistent and elastic registration of histological sections using vector-spline regularization. In Lecture Notes in Computer Science (including subseries Lecture Notes in Artificial Intelligence and Lecture Notes in Bioinformatics).

39. Schmidt, U., Weigert, M., Broaddus, C. and Myers, G. (2018) Cell detection with star-convex polygons. In Lecture Notes in Computer Science (including subseries Lecture Notes in Artificial Intelligence and Lecture Notes in Bioinformatics).

40. Agasti, S.S., Wang, Y., Schueder, F., Sukumar, A., Jungmann, R. and Yin, P. (2017) DNA-barcoded labeling probes for highly multiplexed Exchange-PAINT imaging. Chem. Sci., 10.1039/C6SC05420J.

41. Beechem, J.M. (2020) High-Plex Spatially Resolved RNA and Protein Detection Using Digital Spatial Profiling: A Technology Designed for Immuno-oncology Biomarker Discovery and Translational Research. In Methods in Molecular Biology. pp. 563–583.

42. Schnitzbauer, J., Strauss, M.T., Schlichthaerle, T., Schueder, F. and Jungmann, R. (2017) Super-resolution microscopy with DNA-PAINT. Nat. Protoc., 10.1038/nprot.2017.024.

43. Jungmann, R., Avendaño, M.S., Woehrstein, J.B., Dai, M., Shih, W.M. and Yin, P. (2014) Multiplexed 3D cellular super-resolution imaging with DNA-PAINT and Exchange-PAINT. Nat. Methods, 10.1038/nmeth.2835.

44. Woehrstein, J.B., Strauss, M.T., Ong, L.L., Wei, B., Zhang, D.Y., Jungmann, R. and Yin, P. (2017) Sub-100-nm metafluorophores with digitally tunable optical properties self-assembled from DNA. Sci. Adv., 10.1126/sciadv.1602128.

45. Winkler, J. (2013) Oligonucleotide conjugates for therapeutic applications. Ther. Deliv., 10.4155/tde.13.47.

46. Dovgan, I., Koniev, O., Kolodych, S. and Wagner, A. (2019) Antibody-Oligonucleotide Conjugates as Therapeutic, Imaging, and Detection Agents. Bioconjug. Chem., 10.1021/acs.bioconjchem.9b00306.

47. Stoeckius, M., Hafemeister, C., Stephenson, W., Houck-Loomis, B., Chattopadhyay, P.K., Swerdlow, H., Satija, R. and Smibert, P. (2017) Simultaneous epitope and transcriptome measurement in single cells. Nat. Methods, 10.1038/nmeth.4380.

48. Coulter, A. and Harris, R. (1983) Simplified preparation of rabbit fab fragments. J. Immunol. Methods, 10.1016/0022-1759(83)90031-5.

49. Mariant, M., Camagna, M., Tarditi, L. and Seccamani, E. (1991) A new enzymatic method to obtain high-yield F(ab)2 suitable for clinical use from mouse IgGl. Mol. Immunol., 10.1016/0161-5890(91)90088-2.

50. Sic, H., Kraus, H., Madl, J., Flittner, K.A., Von Münchow, A.L., Pieper, K., Rizzi, M., Kienzler, A.K., Ayata, K., Rauer, S., et al. (2014) Sphingosine-1-phosphate receptors control B-cell migration through signaling components associated with primary immunodeficiencies, chronic lymphocytic leukemia, and multiple sclerosis. J. Allergy Clin. Immunol., 10.1016/j.jaci.2014.01.037.

51. Sarvaria, A., Madrigal, J.A. and Saudemont, A. (2017) B cell regulation in cancer and anti-tumor immunity. Cell. Mol. Immunol., 10.1038/cmi.2017.35.

52. Kuwano, Y., Prazma, C.M., Yazawa, N., Watanabe, R., Ishiura, N., Kumanogoh, A., Okochi, H., Tamaki, K., Fujimoto, M. and Tedder, T.F. (2007) CD83 influences cell-surface MHC class II expression on B cells and other antigen-presenting cells. Int. Immunol., 10.1093/intimm/dxm067.

53. Borges Da Silva, H., Fonseca, R., Pereira, R.M., Cassado, A.A., Álvarez, J.M. and D’Império Lima, M.R. (2015) Splenic macrophage subsets and their function during blood-borne infections. Front. Immunol., 10.3389/fimmu.2015.00480.

54. Schneider, C.A., Rasband, W.S. and Eliceiri, K.W. (2012) NIH Image to ImageJ: 25 years of image analysis. Nat. Methods, 10.1038/nmeth.2089.

55. Walzer, T., Bléry, M., Chaix, J., Fuseri, N., Chasson, L., Robbins, S.H., Jaeger, S., André, P., Gauthier, L., Daniel, L., et al. (2007) Identification, activation, and selective in vivo ablation of mouse NK cells via NKp46. Proc. Natl. Acad. Sci. U. S. A., 10.1073/pnas.0609692104.

56. McCulloch, L., Alfieri, A. and McColl, B.W. (2018) Experimental stroke differentially affects discrete subpopulations of splenic macrophages. Front. Immunol., 10.3389/fimmu.2018.01108.

57. Mebius, R.E. and Kraal, G. (2005) Structure and function of the spleen. Nat. Rev. Immunol., 10.1038/nri1669.

58. Murray, P.J. and Wynn, T.A. (2011) Protective and pathogenic functions of macrophage subsets. Nat. Rev. Immunol., 10.1038/nri3073.

59. Sheng, J., Ruedl, C. and Karjalainen, K. (2015) Most Tissue-Resident Macrophages Except Microglia Are Derived from Fetal Hematopoietic Stem Cells. Immunity, 10.1016/j.immuni.2015.07.016.

60. Martinez-Pomares, L. and Gordon, S. (2012) CD169 + macrophages at the crossroads of antigen presentation. Trends Immunol., 10.1016/j.it.2011.11.001.

61. Veninga, H., Borg, E.G.F., Vreeman, K., Taylor, P.R., Kalay, H., van Kooyk, Y., Kraal, G., Martinez-Pomares, L. and den Haan, J.M.M. (2015) Antigen targeting reveals splenic CD169 + macrophages as promoters of germinal center B-cell responses. Eur. J. Immunol., 10.1002/eji.201444983.

62. Accolla, R.S., Lombardo, L., Abdallah, R., Raval, G., Forlani, G. and Tosi, G. (2014) Boosting the MHC class II-restricted tumor antigen presentation to CD4+ T helper cells: A critical issue for triggering protective immunity and re-orienting the tumor microenvironment toward an anti-tumor state. Front. Oncol., 10.3389/fonc.2014.00032.

63. Murakami, T., Chen, X., Hase, K., Sakamoto, A., Nishigaki, C. and Ohno, H. (2007) Splenic CD19-CD35+B220+ cells function as an inducer of follicular dendritic cell network formation. Blood, 10.1182/blood-2007-01-068387.

64. Singh, N., Avigan, Z.M., Kliegel, J.A., Shuch, B.M., Montgomery, R.R., Moeckel, G.W. and Cantley, L.G. (2019) Development of a 2-dimensional atlas of the human kidney with imaging mass cytometry. JCI Insight, 10.1172/jci.insight.129477.

65. Nielsen, S., Frøkiær, J., Marples, D., Kwon, T.H., Agre, P. and Knepper, M.A. (2002) Aquaporins in the kidney: From molecules to medicine. Physiol. Rev., 10.1152/physrev.00024.2001.

66. Hatta, K., Takagi, S., Fujisawa, H. and Takeichi, M. (1987) Spatial and temporal expression pattern of N-cadherin cell adhesion molecules correlated with morphogenetic processes of chicken embryos. Dev. Biol., 10.1016/0012-1606(87)90119-9.

67. Skinnider, B.F., Folpe, A.L., Hennigar, R.A., Lim, S.D., Cohen, C., Tamboli, P., Young, A., De Peralta-Venturina, M. and Amin, M.B. (2005) Distribution of cytokeratins and vimentin in adult renal neoplasms and normal renal tissue: Potential utility of a cytokeratin antibody panel in the differential diagnosis of renal tumors. Am. J. Surg. Pathol., 10.1097/01.pas.0000163362.78475.63.

68. Schiano, G., Glaudemans, B., Olinger, E., Goelz, N., Müller, M., Loffing-Cueni, D., Deschenes, G., Loffing, J. and Devuyst, O. (2019) The Urinary Excretion of Uromodulin is Regulated by the Potassium Channel ROMK. Sci. Rep., 10.1038/s41598-019-55771-x.

69. Fissell, W.H. and Miner, J.H. (2018) What is the glomerular ultrafiltration barrier? J. Am. Soc. Nephrol., 10.1681/ASN.2018050490.

70. Palmer, R.E., Kotsianti, A., Cadman, B., Boyd, T., Gerald, W. and Haber, D.A. (2001) WT1 regulates the expression of the major glomerular podocyte membrane protein Podocalyxin. Curr. Biol., 10.1016/S0960-9822(01)00560-7.

71. Johnson, R.J., Iida, H., Alpers, C.E., Majesky, M.W., Schwartz, S.M., Pritzl, P., Gordon, K. and Gown, A.M. (1991) Expression of smooth muscle cell phenotype by rat mesangial cells in immune complex nephritis: α-Smooth muscle actin is a marker of mesangial cell proliferation. J. Clin. Invest., 10.1172/JCI115089.

72. Qi, H., Casalena, G., Shi, S., Yu, L., Ebefors, K., Sun, Y., Zhang, W., D’Agati, V., Schlondorff, D., Haraldsson, B., et al. (2017) Glomerular endothelial mitochondrial dysfunction is essential and characteristic of diabetic kidney disease susceptibility. Diabetes, 10.2337/db16-0695.

73. Miner, J.H. (2012) The glomerular basement membrane. Exp. Cell Res., 10.1016/j.yexcr.2012.02.031.

74. Miner, J.H. (1999) Renal basement membrane components. Kidney Int., 10.1046/j.1523-1755.1999.00785.x.

75. Bábíčková, J., Klinkhammer, B.M., Buhl, E.M., Djudjaj, S., Hoss, M., Heymann, F., Tacke, F., Floege, J., Becker, J.U. and Boor, P. (2017) Regardless of etiology, progressive renal disease causes ultrastructural and functional alterations of peritubular capillaries. Kidney Int., 10.1016/j.kint.2016.07.038.

76. Boor, P. and Floege, J. (2012) The renal (myo-)fibroblast: A heterogeneous group of cells. Nephrol. Dial. Transplant., 10.1093/ndt/gfs296.

77. Lely, A.T., Hamming, I., van Goor, H. and Navis, G.J. (2004) Renal ACE2 expression in human kidney disease. J. Pathol., 10.1002/path.1670.

78. Hamming, I., Timens, W., Bulthuis, M.L.C., Lely, A.T., Navis, G.J. and van Goor, H. (2004) Tissue distribution of ACE2 protein, the functional receptor for SARS coronavirus. A first step in understanding SARS pathogenesis. J. Pathol., 10.1002/path.1570.

79. Belliere, J., Casemayou, A., Ducasse, L., Zakaroff-Girard, A., Martins, F., Iacovoni, J.S., Guilbeau-Frugier, C., Buffin-Meyer, B., Pipy, B., Chauveau, D., et al. (2015) Specific macrophage subtypes influence the progression of rhabdomyolysis-induced kidney injury. J. Am. Soc. Nephrol., 10.1681/ASN.2014040320.

80. Ali, M.M., Li, F., Zhang, Z., Zhang, K., Kang, D.K., Ankrum, J.A., Le, X.C. and Zhao, W. (2014) Rolling circle amplification: A versatile tool for chemical biology, materials science and medicine. Chem. Soc. Rev., 10.1039/c3cs60439j.

81. Mullis, K.B. and Faloona, F.A. (1987) Specific Synthesis of DNA in Vitro via a Polymerase-Catalyzed Chain Reaction. Methods Enzymol., 10.1016/0076-6879(87)55023-6.

82. Li, J.J., Chu, Y., Lee, B.Y.H. and Xie, X.S. (2008) Enzymatic signal amplification of molecular beacons for sensitive DNA detection. Nucleic Acids Res., 10.1093/nar/gkn033.

83. Collins, M.L., Irvine, B., Tyner, D., Fine, E., Zayati, C., Chang, C.A., Horn, T., Ahle, D., Detmer, J., Shen, L.P., et al. (1997) A branched DNA signal amplification assay for quantification of nucleic acid targets below 100 molecules/ml. Nucleic Acids Res., 10.1093/nar/25.15.2979.

84. Niekamp, S., Stuurman, N. and Vale, R.D. (2020) A 6-nm ultra-photostable DNA FluoroCube for fluorescence imaging. Nat. Methods, 10.1038/s41592-020-0782-3.

85. Dagogo-Jack, I. and Shaw, A.T. (2018) Tumour heterogeneity and resistance to cancer therapies. Nat. Rev. Clin. Oncol., 10.1038/nrclinonc.2017.166.

86. Johnson, D.B., Bordeaux, J., Kim, J.Y., Vaupel, C., Rimm, D.L., Ho, T.H., Joseph, R.W., Daud, A.I., Conry, R.M., Gaughan, E.M., et al. (2018) Quantitative spatial profiling of PD-1/PD-L1 interaction and HLA-DR/IDO-1 predicts improved outcomes of anti-PD-1 therapies in metastatic melanoma. Clin. Cancer Res., 10.1158/1078-0432.CCR-18-0309.

87. Satija, R., Farrell, J.A., Gennert, D., Schier, A.F. and Regev, A. (2015) Spatial reconstruction of single-cell gene expression data. Nat. Biotechnol., 10.1038/nbt.3192.

88. Halpern, K.B., Shenhav, R., Matcovitch-Natan, O., Tóth, B., Lemze, D., Golan, M., Massasa, E.E., Baydatch, S., Landen, S., Moor, A.E., et al. (2017) Single-cell spatial reconstruction reveals global division of labour in the mammalian liver. Nature, 10.1038/nature21065.

