## Supplementary materials for "SeqStain using fluorescent-DNA conjugated antibodies allows efficient, multiplexed, spatialomic profiling of human and murine tissues"

### SUPPLEMENTARY TABLES, FIGURES AND MOVIES

#### Supplementary Tables

**Table S1**. List of antibodies and Fabs, clones and manufacturers used for immunofluorescence staining and SeqStain modification.

**Table S2.** List of DNA oligonucleotides and their sequences used for SeqStain modification of antibodies and Fabs.

#### Supplementary Figures

**Figure S1**. **Generation of SeqStain antibodies. A**. Schematic showing the layout of the different oligos used for SeqStain modification. **B**. Schematic detailing the steps of antibody modification for SeqStain. In Step1, the antibodies are conjugated to the linker oligo using various conjugation chemistries (methods). In Step 2, the rest of the oligo complex containing the docking oligo and the fluorescent oligo is annealed to the linker oligo that is conjugated to the antibody. **C**. Image of SDS-PAGE gel showing analysis of antibody conjugation to linker oligos (Step 1). Arrows show the bands representing the conjugated antibody heavy and light chains compared and the unmodified antibodies. The different bands correspond to the differences in the number of conjugated linker oligo per heavy chain. **D.** Image of agarose gel showing analysis of conjugated and annealed SeqStain antibodies. The annealed SeqStain-ready antibodies can be seen as a shifted band compared to the unbound oligo complex. Arrows show the bands representing the fluorescent DNA-hybridized antibodies and the unbound DNA complex.

**Figure S2**. **Staining with SeqStain antibodies. A.** Flow cytometric analysis of RAW264.7 cells stained with anti-CD45 and anti-CD11b SeqStain and conventional antibodies. Cells stained with fluorescent oligo alone and unstained cells were used as control. **B.** Representative immunofluorescence images of immobilized RAW 264.7 cells stained with CD45 SeqStain antibody or by the conventional immunostaining method. Cells stained with fluorescent oligo alone was used as a control. The antibodies were labelled using AF488 fluorophore (shown in green). Scale bar is 100μm. **C.** Representative immunofluorescent images showing RAW 264.7 cells stained antibody conjugated with linker oligo alone (without hybridization with fluorescent-DNA complex) (Left panel) or stained with the corresponding unmodified antibody (Right panel). The antibodies were co-stained with AF488 containing anti-rat secondary antibody. Scale bar is 100μm.

**Figure S3. De-staining time course in SeqStain.** Immunofluorescence images showing RAW264.7 cells stained with anti-CD45 SeqStain antibody, de-stained with DNase I and imaged at various time points. Images were acquired at different time points post DNase I addition as indicated in each panel. A bar graph showing quantification of fluorescence intensity after staining (red bar) and de-staining (brown bars) in each panel is also presented (bottom). Graphs show the mean ± standard deviation. Scale bar is 100μm.

**Figure S4. Generation of SeqStain antibodies using DBCO-Azide and biotin-streptavidin method. A and B.** Schematics showing preparation of SeqStain antibodies using (**A**) DBCO-Azide click chemistry or the (**B**) biotin-streptavidin chemistry to conjugate linker oligos to antibodies. **C.** Representative immunofluorescence images of K562 human myelogenous leukemia cells stably expressing CD11b and CD18 proteins and stained with anti-CD11b SeqStain antibody and de-stained with DNase I. The anti-CD11b antibody was conjugated to the linker oligo using DBCO-Azide click chemistry and subsequently hybridized to complementary fluorescent-DNA complex. The antibodies were labelled with AF488 fluorophore (shown in green). Scale bar is 100μm. A graph showing quantification of fluorescence intensity after staining (green bars) and de-staining (brown bars) in each panel is also presented. Graphs show the mean ± standard deviation. **D.** Representative immunofluorescence image of RAW264.7 cells stained with anti-CD11b SeqStain antibody and de-stained with DNase I. Scale bar is 100μm. The CD11b antibody was conjugated to the linker oligo using the biotin-streptavidin chemistry and subsequently hybridized to complementary fluorescent-DNA complex. The antibodies were labelled with AF488 fluorophore (shown in green). A graph showing quantification of fluorescence intensity after staining (green bars) and de-staining (brown bars) in each panel is also presented. Graphs show the mean ± standard deviation.

**Figure S5. Repeat staining and de-staining of RAW264.7 cells with the same set of SeqStain antibodies.** Immunofluorescence images of RAW264.7 cells repeatedly co-stained in the first five cycles with anti-CD44 SeqStain antibody bearing AF488 fluorophore and anti-CD45 SeqStain antibody bearing Cy3 fluorophore (left panel) and de-stained using DNase I (right panel). Subsequently, the cells were repeatedly stained with anti-CD68 SeqStain antibody bearing AF488 fluorophore and anti-CD11b SeqStain antibody bearing Cy3 fluorophore in the next five rounds. A graph showing quantification of fluorescence intensity after staining (green and red bars) and de-staining (brown bars) in each panel is also presented. Graphs show the mean ± standard deviation. Scale bar is 100μm.

**Figure S6. Generation of SeqStain Fabs**. **A.** Schematic showing preparation of SeqStain Fabs. In step1, the Fab fragments are conjugated to the linker oligos. In step 2, the modified Fab is hybridized to fluorescent-DNA complex. **B**. Image of SDS-PAGE gel showing analysis of Fab conjugation to linker oligos (Step 1). The different bands correspond to the differences in the number of conjugated linker oligo per Fab. **C.** Image of agarose gel showing analysis of Fabs after the annealing step (Step 2). Arrows show the bands representing the fluorescent DNA-hybridized Fabs and the unbound DNA complex.

**Figure S7. Multiplex staining of RAW264.7 cells using SeqStain Fabs:** Immunofluorescence images of RAW264.7 cells after each of the three rounds of staining with two unique antibodies pre-complexed with two different SeqStain Fabs (with Fabs labelled using the AF488 fluorophore shown in green and the Fabs labelled using the AF546 fluorophore shown in red) and after de-staining with DNase I. The primary antibodies used in each round are indicated in the panel. All images are representative of at least three replicates and different fields from each round are presented here to show representation. Scale bar is 100$\mu$m. A graph showing quantification of the fluorescence intensity after staining (green and red bars) and de-staining (brown bars) in each panel is also presented on the right. Graphs show mean $\pm$ standard deviation.

**Figure S8: Staining tissues with SeqStain antibodies labelled with different fluorophores.** Representative immunofluorescence images of human kidney tissue sections stained with anti-CD31, anti-Cytokeratin-8 or anti-Collagen-IV SeqStain antibodies labelled with either AF488 fluorophore (Left panels) or a spectrally different fluorophore (as labelled, Right panels). All images are representative of at least three replicates. Scale bar is 100$\mu$m.

**Figure S9: Only SeqStain antibodies are de-stained upon treatment with a nuclease. A**. Representative immunofluorescence images of murine tumor tissues (LLC tumors) after staining with DAPI (blue) and the anti-CD45 antibody (green) using either the SeqStain technique (top panel) or the conventional immunostaining methodology (bottom panel). Representative images post-destaining with DNase I are also presented (Right panels). The antibodies were labelled using the AF488 fluorophore. Scale bar is 100$\mu$m. A bar graph showing quantification of fluorescence intensity after staining (green and blue bars) and de-staining (brown bars) in each panel is also presented (right). Graph shows mean ± standard deviation. **B**. Representative immunofluorescence images of human kidney tissues after staining with DAPI (blue) and the anti-Nephrin antibody either pre-complexed with SeqStain Fab (top panel) or the conventional immunostaining method (bottom panel). Representative images acquired after DNase I treatment are also shown (right panels). Scale bar is 100$\mu$m. A graph showing quantification of fluorescence intensity after staining (green and blue bars) and de-staining (brown bars) in each panel is also presented. Graph shows the mean ± standard deviation.

**Figure S10: SeqStain de-staining step using DNase I treatment does not affect the nuclear DNA proteins or the DNA.** Representative immunofluorescence images of human kidney sections treated with DNase I either once (1X) or repeatedly (5X, 10X) and stained for a common DNA binding protein (Histone H1) by conventional immunofluorescence methodology. Top panels show Histone H1 immunofluorescence staining in each of the conditions and the bottom panels show DAPI staining. All images are representative of at least three replicates. Scale bar is 100μm. A bar graph showing quantification of the fluorescence intensity (green and blue bars) is also presented on the right. Graphs show mean ± standard deviation.

**Figure S11: SeqStain methodology allows selective removal of fluorophores linked to SeqStain antibodies without affecting fluorophores on conventional, commercial antibodies. A.** Representative immunofluorescence images of human kidney tissue sections stained with anti-Synaptopodin SeqStain antibody (AF488, green) and counter-stained with corresponding secondary antibody (AF546, red). The individual channel images and the merged image are shown in individual panels (top). Representative images acquired after DNase I treatment are shown in the bottom panels. Scale bar is 100μm. A graph showing quantification of fluorescence intensity after staining (green and red bars) and de-staining (brown bars) in each panel is also presented. Graphs show the mean ± standard deviation. **B**. Representative immunofluorescence staining of human kidney tissue with anti-CD68 SeqStain antibody (AF488, green) and counter-stained with corresponding secondary antibody (AF546, red). The individual channel images and the merged image are shown in individual panels (top). Representative images acquired after DNase I treatment are shown in the bottom panels. Scale bar is 100μm. A graph showing quantification of fluorescence intensity after staining (green and red bars) and de-staining (brown bars) in each panel is also presented. Graphs show the mean ± standard deviation.

**Figure S12: SeqStain secondary antibodies provide staining pattern on complex tissues similar to conventional immunofluorescence staining.** Representative immunofluorescence images of human kidney tissue section stained with anti-EpCAM or anti-Collagen antibodies pre-complexed with SeqStain secondary antibodies (top panels) and de-stained using DNase I ( bottom panels). Secondary antibodies bearing AF488 fluorophore are shown in green and with Cy3 fluorophore are shown in red. All images are representative of at least three replicates. Scale bar is 100μm. A graph showing quantification of the fluorescence intensity after staining (green and red bars) and de-staining (brown bars) in each panel is also presented on the right. Graphs show mean + standard deviation.

**Figure S13**: **SeqStain based multiplexed staining and de-staining of murine spleen tissue.** Whole slide images of murine spleen tissue acquired after each of the five cycles of staining with unique SeqStain antibodies and de-staining with DNase I. The antibodies used in each round are indicated in the panel, with SeqStain antibodies labelled using the AF488 fluorophore shown in green and the antibodies labelled using the AF546 fluorophore shown in red. Scale bar is 100μm.

**Figure S14: Cell segmentation of SeqStain stained tissue sections using Cell Profiler and Fiji. A**. Immunofluorescent images (Left panels) and the corresponding Cell Profiler analysed images (Right panels) of mouse spleen tissue stained with CD11b SeqStain antibody. DAPI stained nuclei were demarcated and identified as primary objects (Top panels) while cells stained with CD11b was demarcated and identified as secondary objects (middle panel). The primary and secondary objects were linked and quantified using Cell Profiler (Bottom panel). **B.** Immunofluorescent images (Left panels) and the corresponding Cell Profiler analysed images (Right panels) of human kidney tissue stained with CD31 SeqStain antibody. DAPI stained nuclei were demarcated and identified as primary objects (Top panels) while cells stained with CD31 was demarcated and identified as secondary objects (middle panel). The primary and secondary objects were linked and quantified using Cell Profiler (Bottom panel). **C**. Immunofluorescent image of DAPI stained nuclei from a representative region in the multiplex stained spleen tissue (Left panel) and the corresponding StarDist generated segmented nuclei (Right panel).

**Figure S15. SeqStain-based 25-plex staining and imaging of whole human kidney tissue.** Immunofluorescence images of whole kidney tissue sections stained with SeqStain antibodies and DAPI (as indicated in the panel) in 25-plex experiment. Antibody used in each staining step is labelled on the panel. Zoomed-in sections from each of the immunofluorescence images are presented below each panel. Images are representative of at least two replicates. Scale bar is 100μm.

**Figure S16: Signal integrity of the staining obtained with SeqStain antibodies.** The fluorescence intensity profile was compared for two of the markers (Vimentin and AQP1) that were repeated during the multiplex staining of human kidney. The intensity profile was measured by ImageJ plot profiler around the indicated yellow line. The cycle number is as indicated in the panel.

**Supplementary Movies**

**Supplementary Movie 1. Time-lapse of video showing the rate of de-staining on RAW 264.7 cells.** RAW264.7 cells were co-stained with anti-CD44 SeqStain AF488 and anti-CD45 SeqStain Cy3 antibodies. Subsequently, the stained cells were incubated with DNase I for de-staining and the fluorescence images were acquired as a video. Images were acquired every 3 seconds.

**Supplementary Movie 2. Time lapse video showing the rate of de-staining of Human kidney tissues.** Normal human kidney tissue sections were stained with **anti-Cytokeratin 8 SeqStain AF488** antibody. Subsequently, the stained tissue section was incubated with DNase I for de-staining and the fluorescence images were acquired as a video. Images were acquired every 20 seconds.

**Supplementary Movie 3. Time lapse video showing the rate of de-staining of Human kidney tissues.** Normal human kidney tissue sections were stained with anti-Collagen IV SeqStain Cy3 antibody. Subsequently, the stained tissue section was incubated with DNase I for de-staining and the fluorescence images were acquired as a video. Images were acquired every 20 seconds.

**Supplementary Movie 4. Time lapse video showing the rate of de-staining of Human kidney tissues.** Normal human kidney tissue sections were stained with **anti-EpCAM SeqStain Cy5** antibody. Subsequently, the stained tissue section was incubated with DNase I for de-staining and the fluorescence images were acquired as a video. Images were acquired every 20 seconds.

**Table S1.** List of antibodies and Fabs, clones and manufacturers used for immunofluorescence staining and SeqStain modification.

| Antigen | Clone | Source | Catalog number |
|---|---|---|---|
| **Primary antibodies** | | | |
| CD11b | M1/70 | Biolegend | 101248 |
| a-Tubulin | Polyclonal | Abcam | ab18251 |
| CD45 (mouse) | 30-F11 | Biolegend | 103164 |
| Vinculin | EPR8185 | Abcam | ab129002 |
| F4/80 | T45-2342 | BD Biosciences | 565409 |
| Paxillin | Y113 | Abcam | ab32084 |
| CD45R/B220 | RA3-6B2 | BD Biosciences | 557390 |
| CD44 | IM7 | BD Biosciences | 553131 |
| MHC II | M5/114.15.2 | BD Biosciences | 556999 |
| IgD | 11-26c.2a | BD Biosciences | 553438 |
| IgM | R6-60.2 | BD Biosciences | 553405 |
| CD169 | 3D6.112 | Biolegend | 142402 |
| CD34 | QBEND/10 | Thermo Fisher Scientific | MA1-10202 |
| CD68 (mouse) | Y1/82A | Biolegend | 333802 |
| CD45 (human) | H130 | BD Biosciences | 555480 |
| CD68 (human) | KP1 | Thermo Fisher Scientific | 14-0688-80 |
| CD8 | HIT8a | BD Biosciences | 555631 |
| CD31 | WM59 | BD Biosciences | 555444 |
| Cytokeratin-8 | LP3K | eBioscience | 14-9938-82 |
| Cytokeratin-7 | RCK105 | Santa Cruz Biotechnology | sc-23876 |
| Collagen IV | 1042 | Thermo Fisher Scientific | 14-9871-82 |
| Histone | AE-4 | Santa Cruz Biotechnology | sc-8030 |
| Podocin | Polyclonal | Sigma-Aldrich | P0372 |
| Synaptopodin | D9 | Santa Cruz Biotechnology | sc-515842 |
| EpCAM | HEA125 | Santa Cruz Biotechnolgy | sc-59906 |
| Aquaporin 1 | Polyclonal<br><br>B-11 | Millipore Sigma<br><br>Santa Cruz Biotechnology | SAB4501545<br><br>sc-25287 |
| Aquaporin 2 | E-2 | Santa Cruz Biotechnology | sc-515770 |
| Aquaporin 3 | F-1 | Santa Cruz Biotechnology | sc-518001 |
| ACE2 | E-11 | Santa Cruz Biotechnology | sc-390851 |
| WT1 | 6F-H2 | Thermo Fisher Scientific | MA1-46028 |
| Nephrin | Polyclonal | Thermo Fisher Scientific | PA5-72826 |
| Megalin | H-10 | Santa Cruz Biotechnology | sc-515772 |

| | | | |
|---|---|---|---|
| Na+K+-ATPase | C464.6 | Santa Cruz Biotechnology | sc-21712 |
| Uromodulin | 877914 | R&D systems | MAB5144 |
| Vimentin | V9 | Santa Cruz Biotechnology | sc-6260 |
| α-SMA | 1A4 | Millipore Sigma | A5228 |
| **Secondary antibodies** | | | |
| Mouse IgG | Polyclonal | Thermo Fisher Scientific | A21202, A11031 |
| Rabbit IgG | Polyclonal | Thermo Fisher Scientific | A10042, A21206 |
| Rat IgG | Polyclonal | Thermo Fisher Scientific | A11081, A11006 |
| **Fabs** | | | |
| Rabbit IgG, Fc fragment | Polyclonal | Jackson ImmunoResearch | 111-007-008 |
| Mouse IgG1, Fc fragment | Polyclonal | Jackson ImmunoResearch | 115-007-185 |
| Rat IgG, Fcy fragment specific | Polyclonal | Jackson ImmunoResearch | 112-007-008 |

**Table S2.** List of DNA oligonucleotides and their sequences used for SeqStain modification of antibodies and Fabs.

| ID | Sequence | Modification |
|---|---|---|
| mAb linker with EcoRV site | 5'-TTTTTTTTTTAGCAGATATCACAGC | 5'- Amine |
| mAb linker with SmaI site | 5'-TTTTTTTTTTAGCACCCGGGACAGC | 5'-Amine |
| mAb linker with Azide | 5'-ACGGGATATCAGATACGGGATATCAGATACGGGA-TATCAGAT | 3'-Azide |
| mAb linker with Biotin | 5'-ACGGGATATCAGATACGGGATATCAGATACGGGA-TATCAGAT | 3'-Biotin |
| Bridging oligo with EcoRV site | 5'-TTGACAGCTGCCGGATTGACAGCTGCCGGATTGA-CAGCTGCCGGATTGACAGCTGCCGGA-TTGACAGCTGCCGGA GCTGTGATATCTGCT | None |
| Bridging oligo with SmaI site | 5'-TTGACAGCTGCCGGATGACAGCTGCCGGA-TTGACAGCTGCCGGATTGACAGCTGCCGGA-TTGACAGCTGCCGGAGCTGTCCCGGGTGCT | None |
| Fluorescent oligo (Maleimide-Sulfhydryl chemistry) | 5'-TCCGGCAGCTGTCAA | 3' AF488, 3' AF546, 3'-AF594, 3'cy3 |
| Fluorescent oligo (DBCO-Azide chemistry) | 5'-ATCTGATATCCCGT | 3' AF488 |
| Fluorescent oligo (Biotin-Streptavidin chemistry) | 5'-ATCTGATATCCCGT | 3' AF488 |
| Single stranded blocking oligo | 5'-TTTTCCCTCTTCTCTTCCTT | None |

A

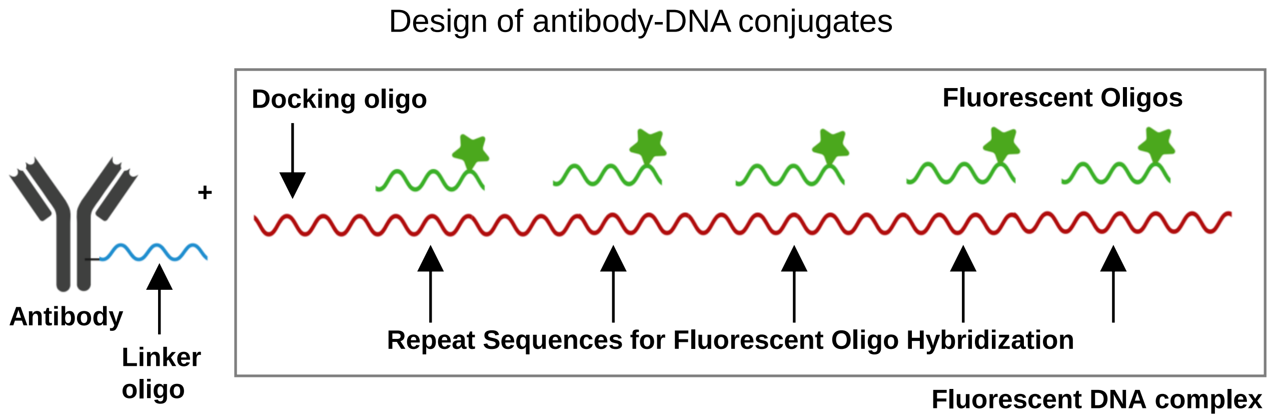


B

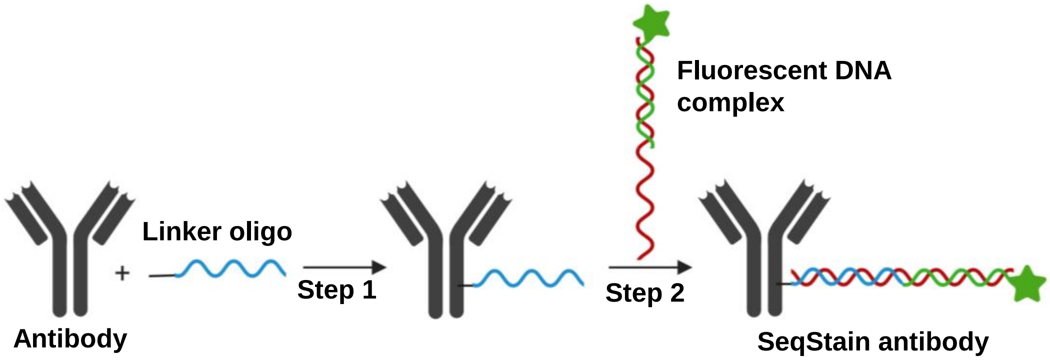


C

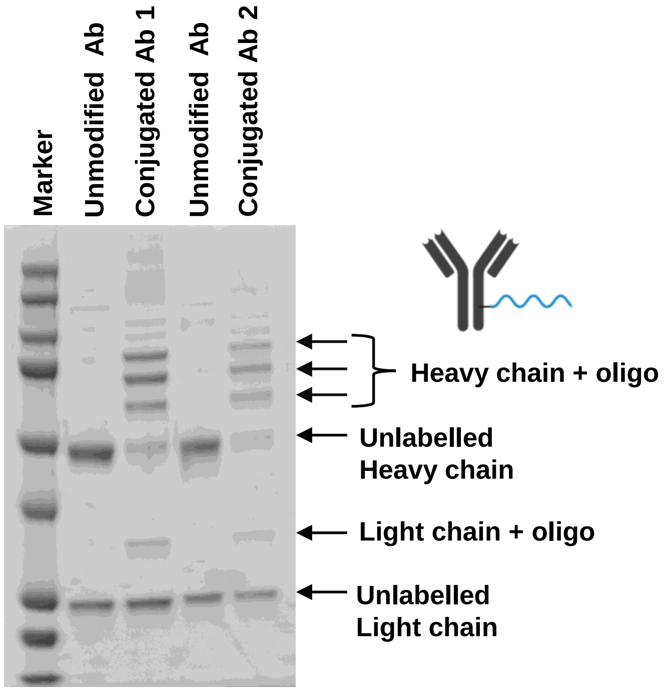


D

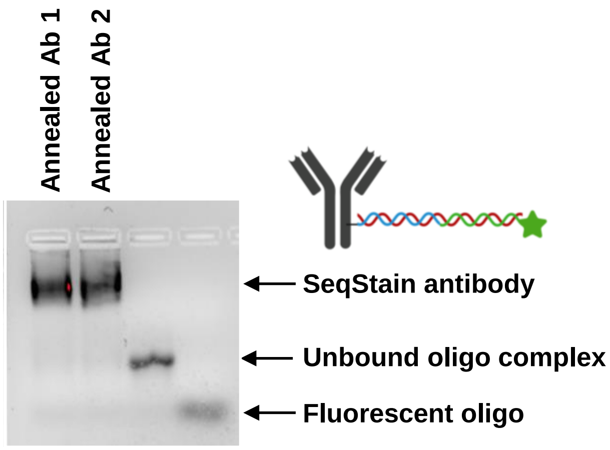


Figure S1

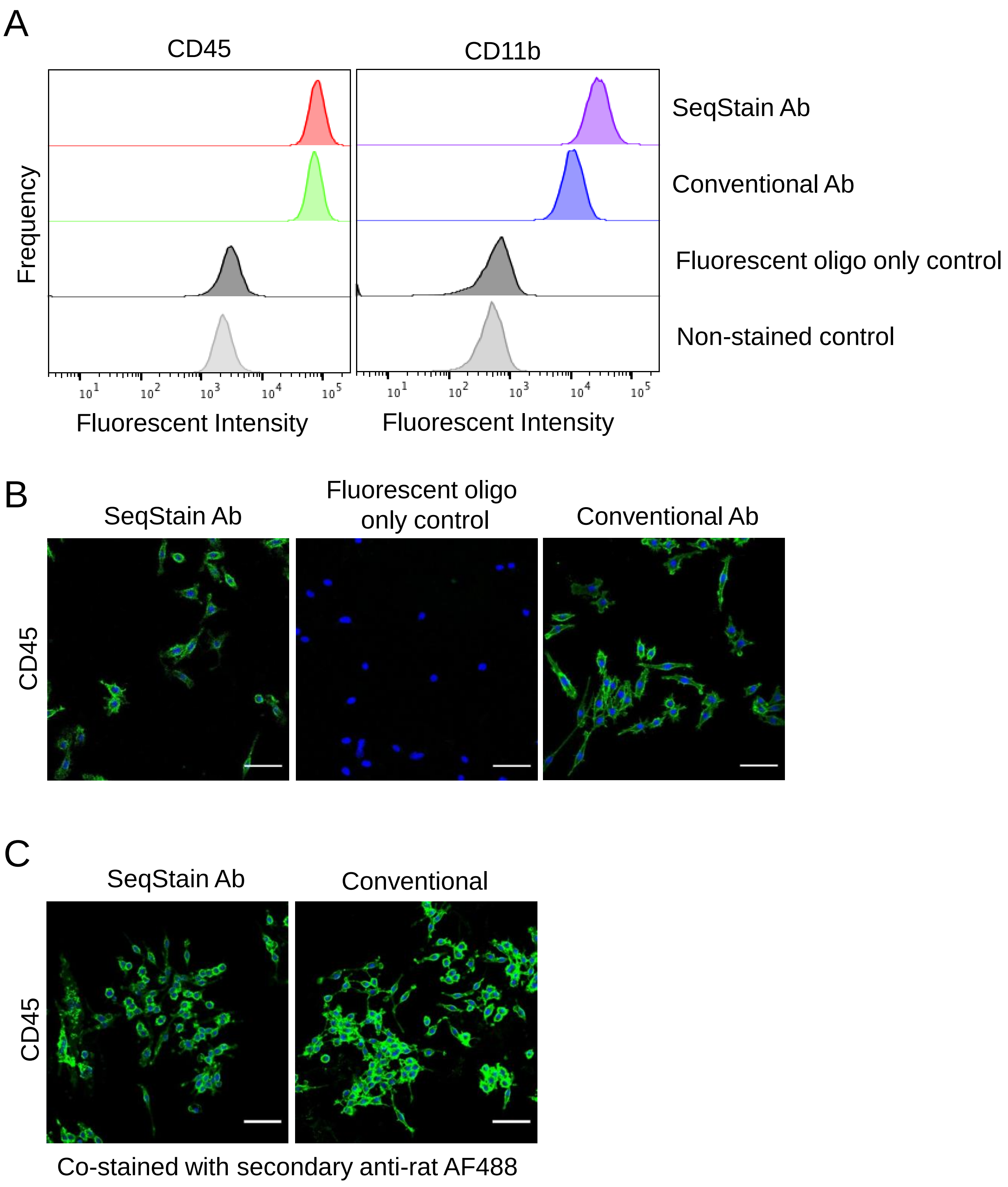
A
CD45
CD11b
Frequency
SeqStain Ab
Conventional Ab
Fluorescent oligo only control
Non-stained control
Fluorescent Intensity
Fluorescent Intensity
B
SeqStain Ab
Fluorescent oligo only control
Conventional Ab
CD45
C
SeqStain Ab
Conventional
CD45
Co-stained with secondary anti-rat AF488

Figure S2

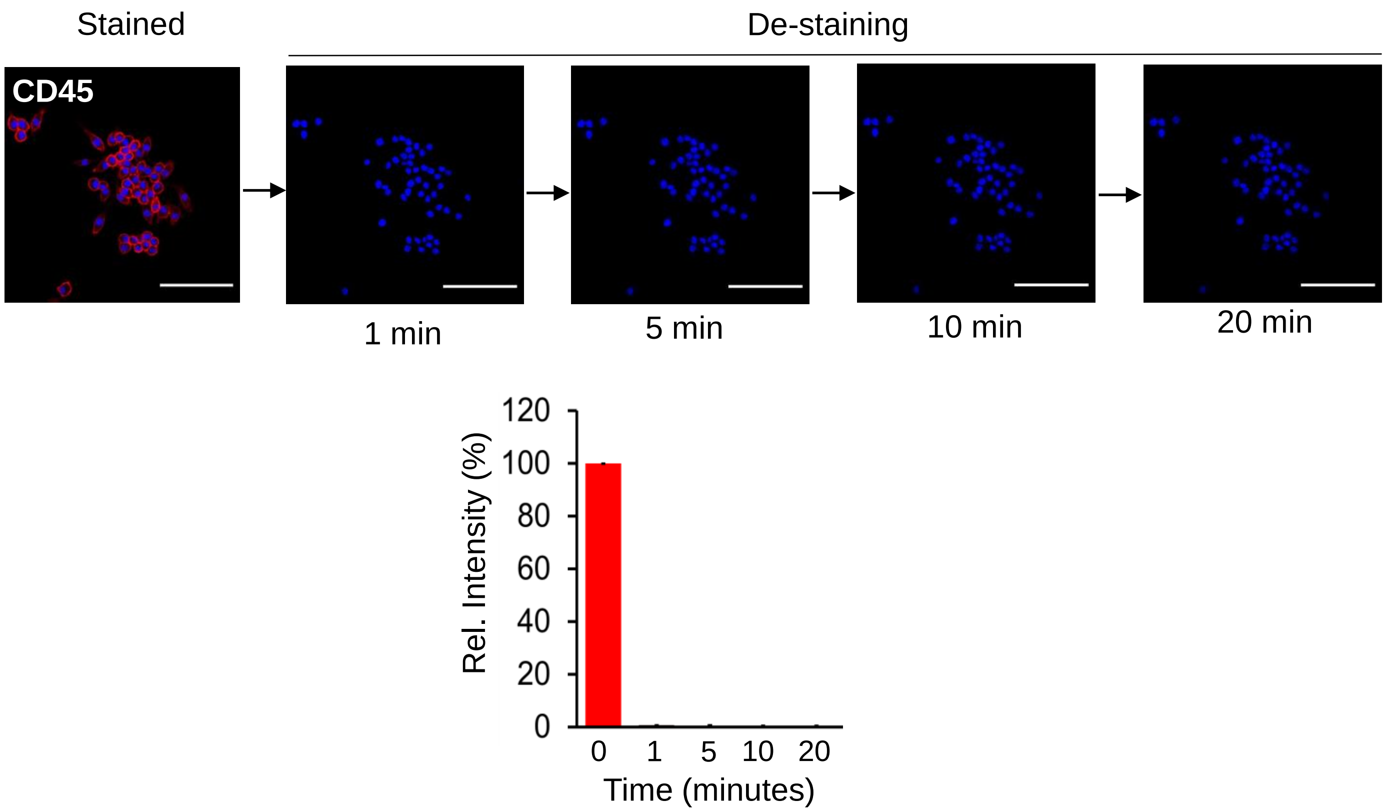
Stained
De-staining
CD45
1 min
5 min
10 min
20 min
Rel. Intensity (%)
120
100
80
60
40
20
0
0
1
5
10
20
Time (minutes)

Figure S3

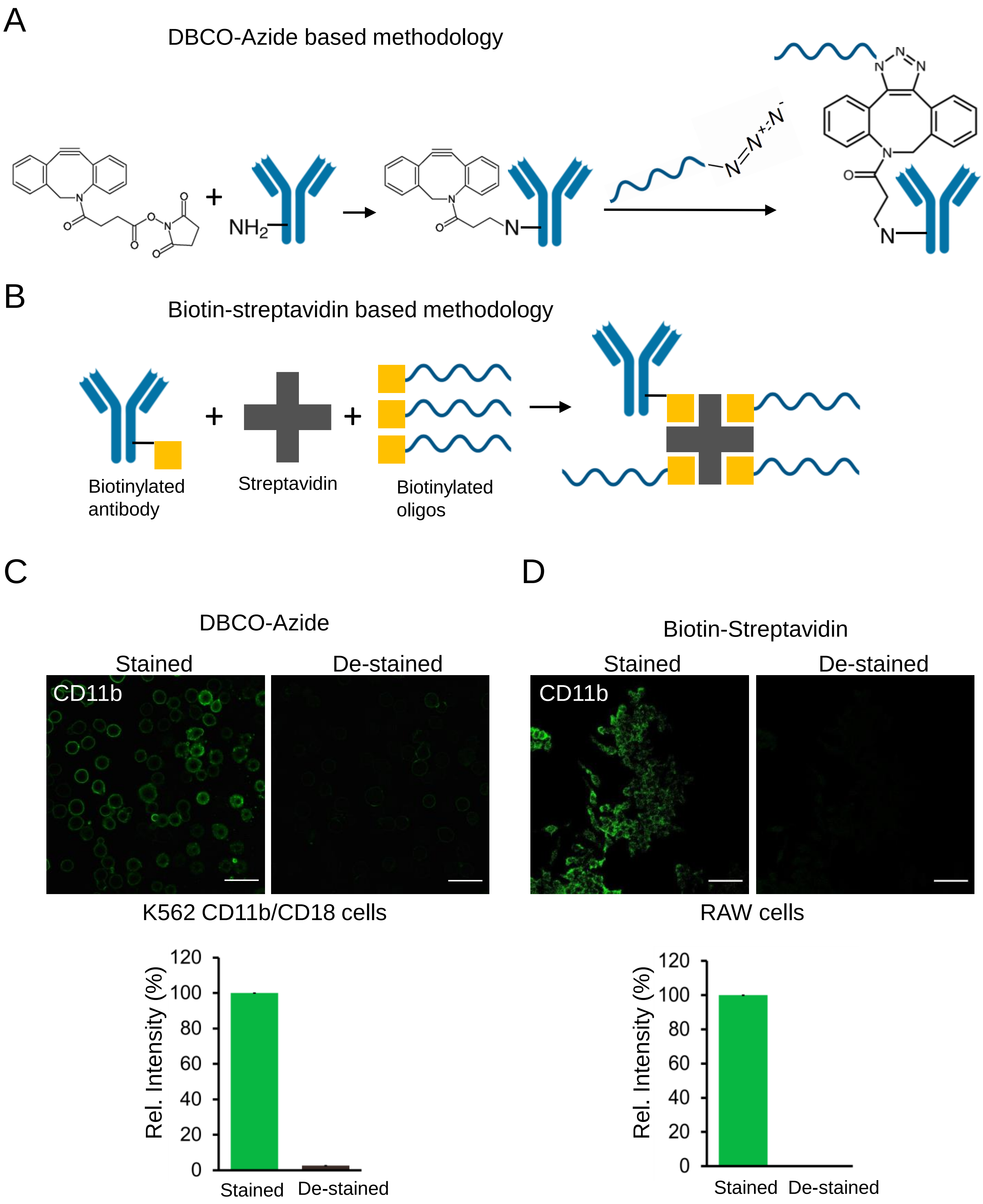
A
DBCO-Azide based methodology
NH2
N
B
Biotin-streptavidin based methodology
Biotinylated antibody
Streptavidin
Biotinylated oligos
C
DBCO-Azide
Stained
De-stained
CD11b
K562 CD11b/CD18 cells
Rel. Intensity (%)
120
100
80
60
40
20
0
Stained
De-stained
D
Biotin-Streptavidin
Stained
De-stained
CD11b
RAW cells
Rel. Intensity (%)
Stained
De-stained

Figure S4

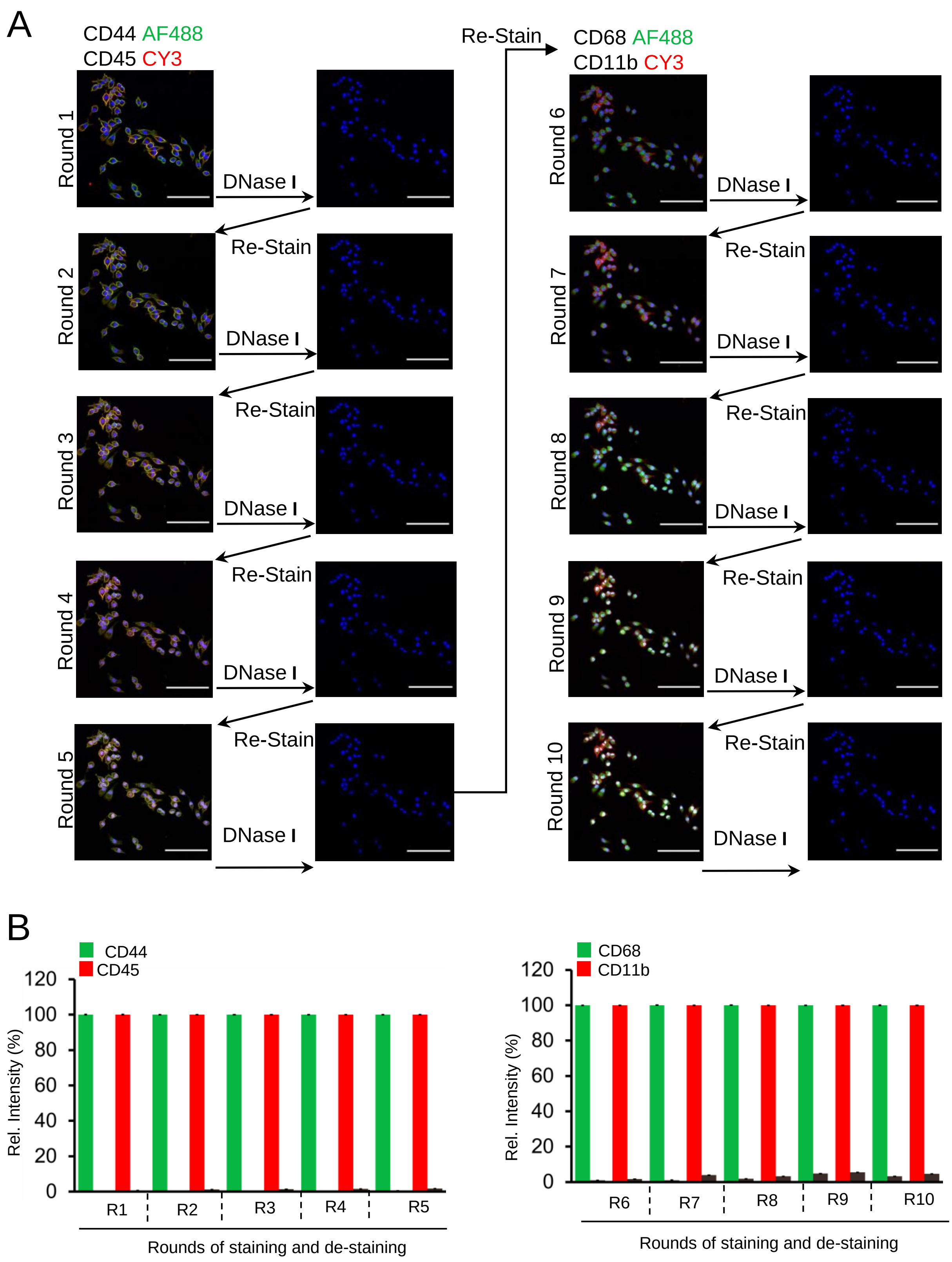

A
CD44 AF488
CD45 CY3
Round 1
Round 2
Round 3
Round 4
Round 5
DNase I
Re-Stain
CD68 AF488
CD11b CY3
Round 6
Round 7
Round 8
Round 9
Round 10
B
CD44
CD45
Rel. Intensity (%)
120
100
80
60
40
20
0
R1
R2
R3
R4
R5
Rounds of staining and de-staining
CD68
CD11b
R6
R7
R8
R9
R10


Figure S5

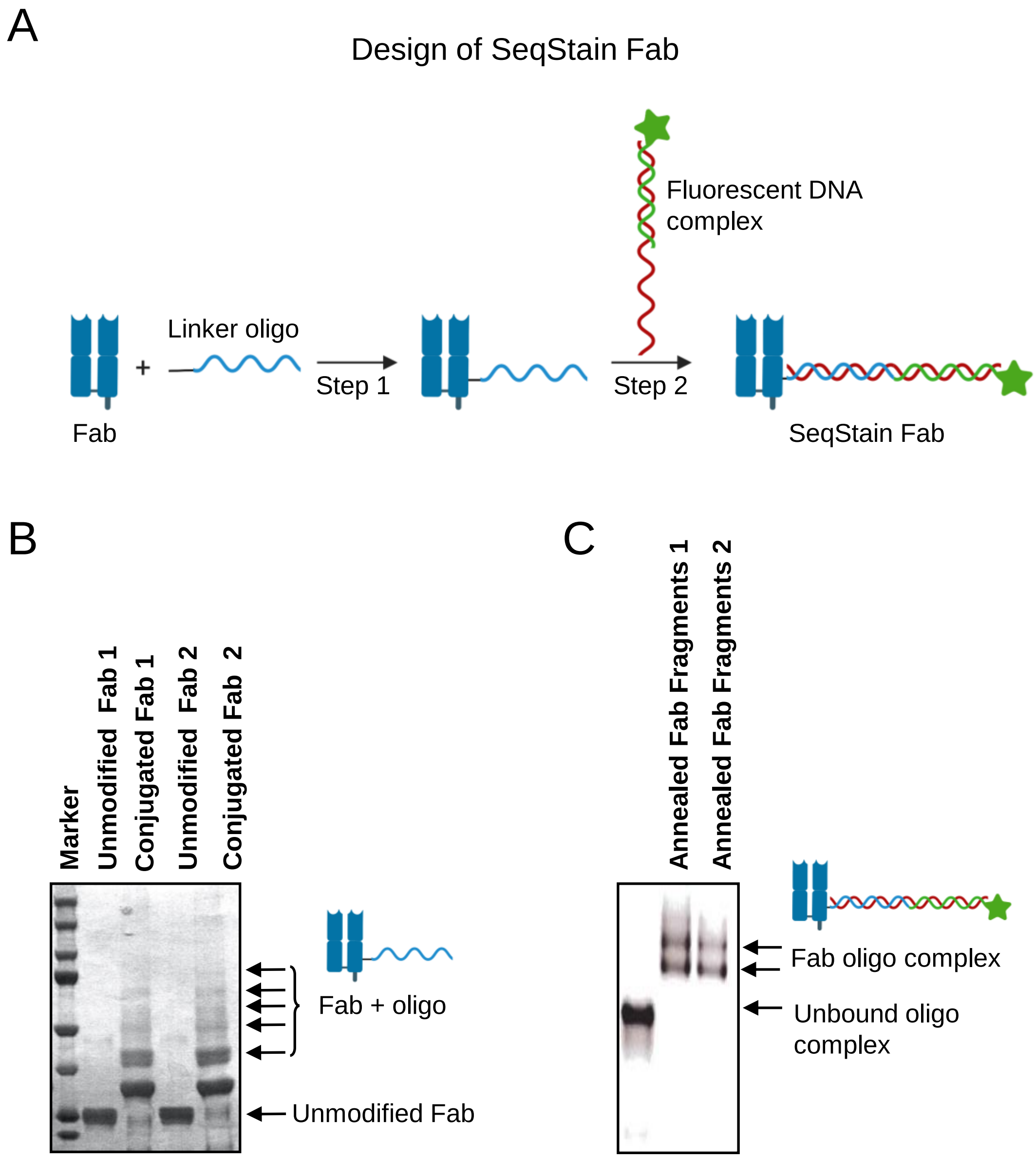
A
Design of SeqStain Fab
Fluorescent DNA
complex
Linker oligo
+
Step 1
Step 2
Fab
SeqStain Fab
B
Marker
Unmodified Fab 1
Conjugated Fab 1
Unmodified Fab 2
Conjugated Fab 2
Fab + oligo
Unmodified Fab
C
Annealed Fab Fragments 1
Annealed Fab Fragments 2
Fab oligo complex
Unbound oligo
complex

Figure S6

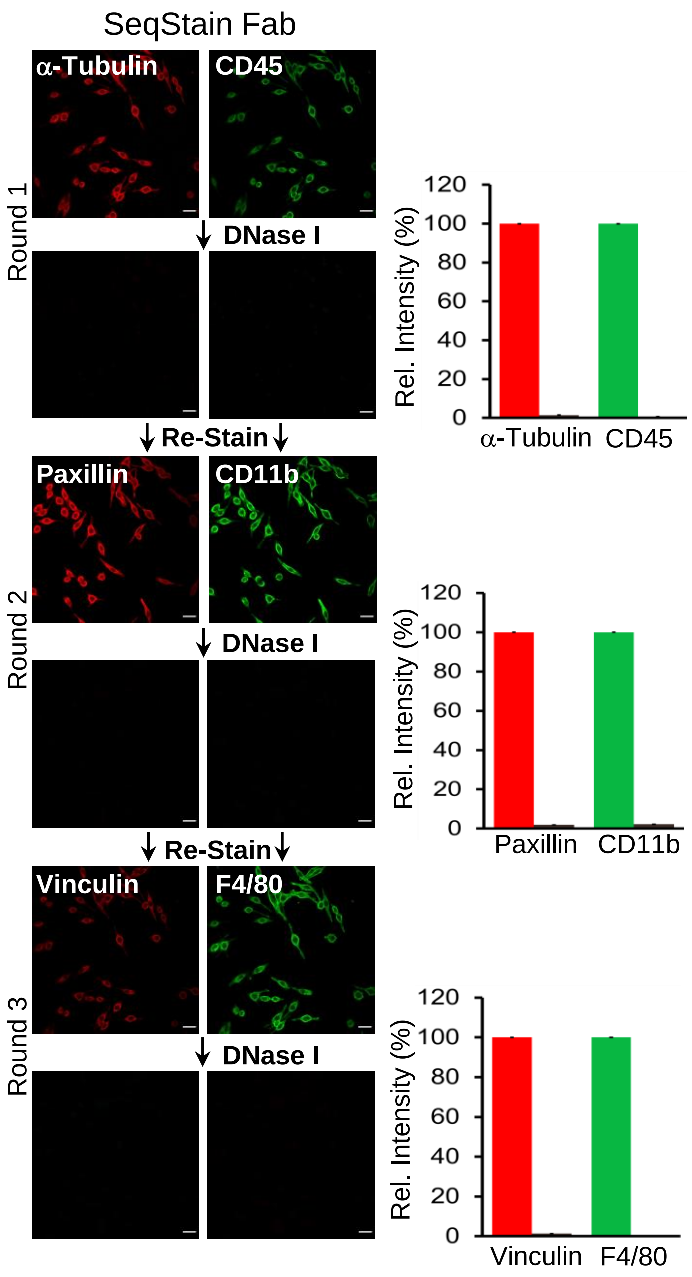

SeqStain Fab
α-Tubulin
CD45
Round 1
DNase I
Re-Stain
Paxillin
CD11b
Round 2
DNase I
Re-Stain
Vinculin
F4/80
Round 3
DNase I
Rel. Intensity (%)
120
100
80
60
40
20
0
α-Tubulin
CD45
Paxillin
CD11b
Vinculin
F4/80


Figure S7

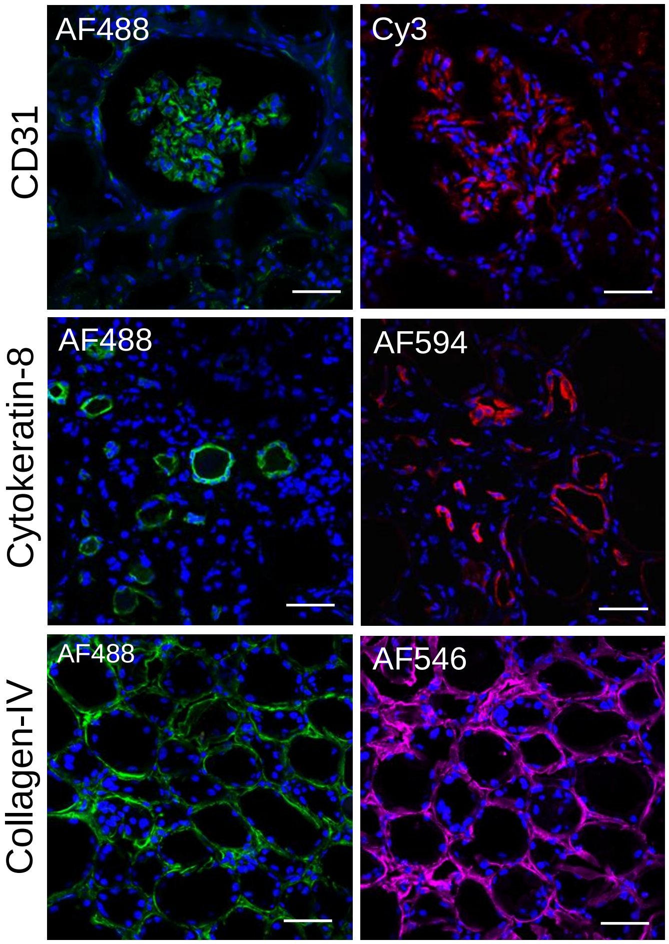
AF488
Cy3
CD31
AF488
AF594
Cytokeratin-8
AF488
AF546
Collagen-IV

Figure S8

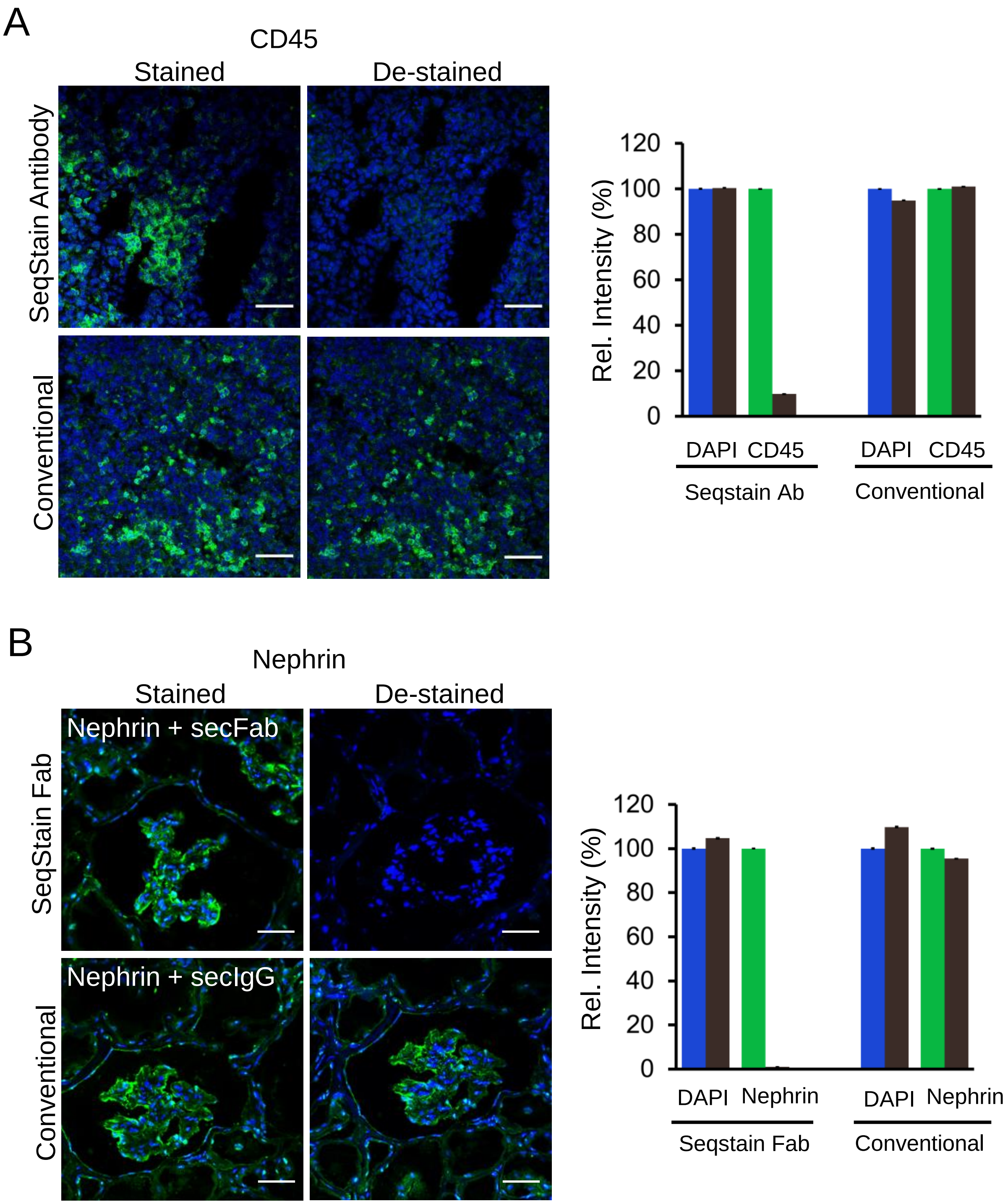
A
CD45
Stained
De-stained
SeqStain Antibody
Conventional
Rel. Intensity (%)
120
100
80
60
40
20
0
DAPI
CD45
DAPI
CD45
Seqstain Ab
Conventional
B
Nephrin
Stained
De-stained
Nephrin + secFab
Nephrin + secIgG
SeqStain Fab
Conventional
Rel. Intensity (%)
120
100
80
60
40
20
0
DAPI
Nephrin
DAPI
Nephrin
Seqstain Fab
Conventional

Figure S9

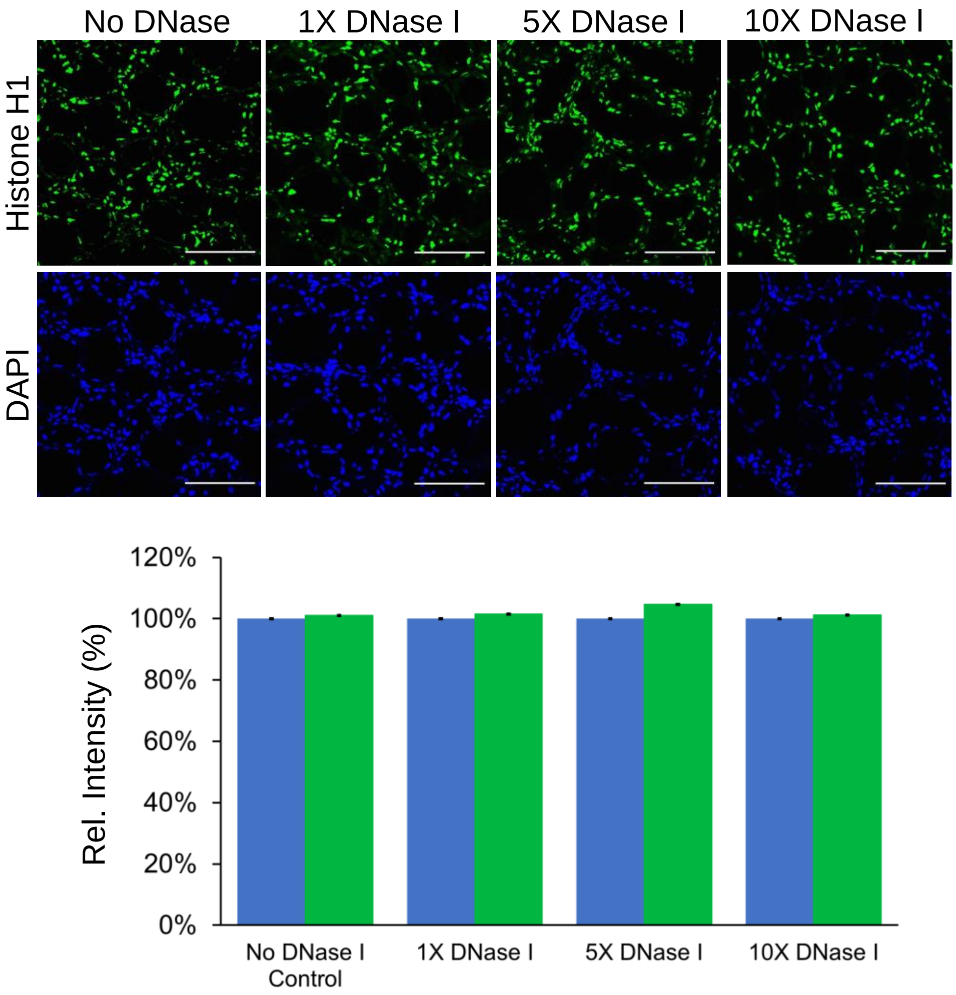
No DNase
1X DNase I
5X DNase I
10X DNase I
Histone H1
DAPI
120%
100%
80%
60%
40%
20%
0%
Rel. Intensity (%)
No DNase I Control
1X DNase I
5X DNase I
10X DNase I

Figure S10

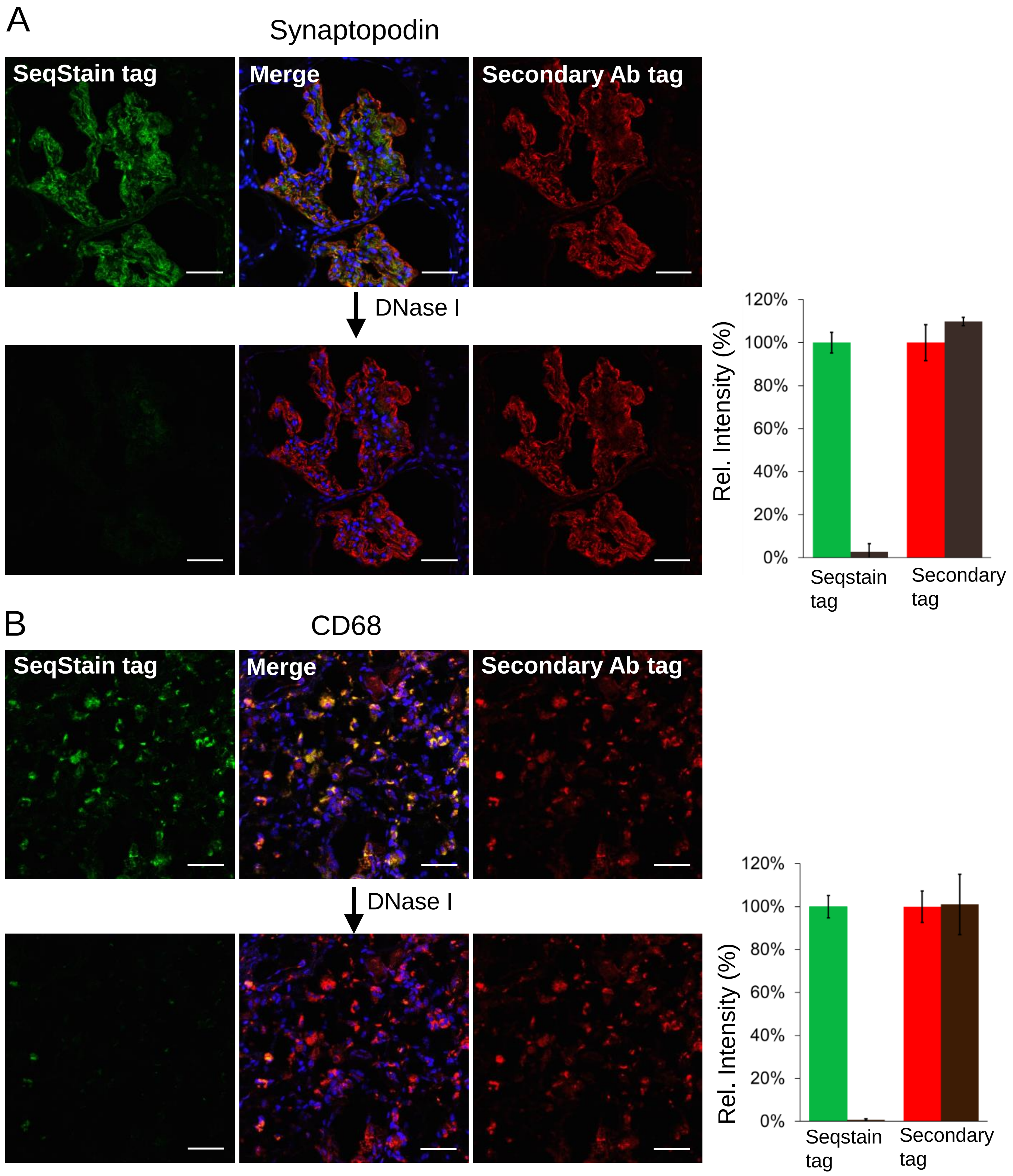
A
Synaptopodin
SeqStain tag
Merge
Secondary Ab tag
DNase I
120%
100%
80%
60%
40%
20%
0%
Rel. Intensity (%)
Seqstain tag
Secondary tag
B
CD68
SeqStain tag
Merge
Secondary Ab tag
DNase I
120%
100%
80%
60%
40%
20%
0%
Rel. Intensity (%)
Seqstain tag
Secondary tag

Figure S11

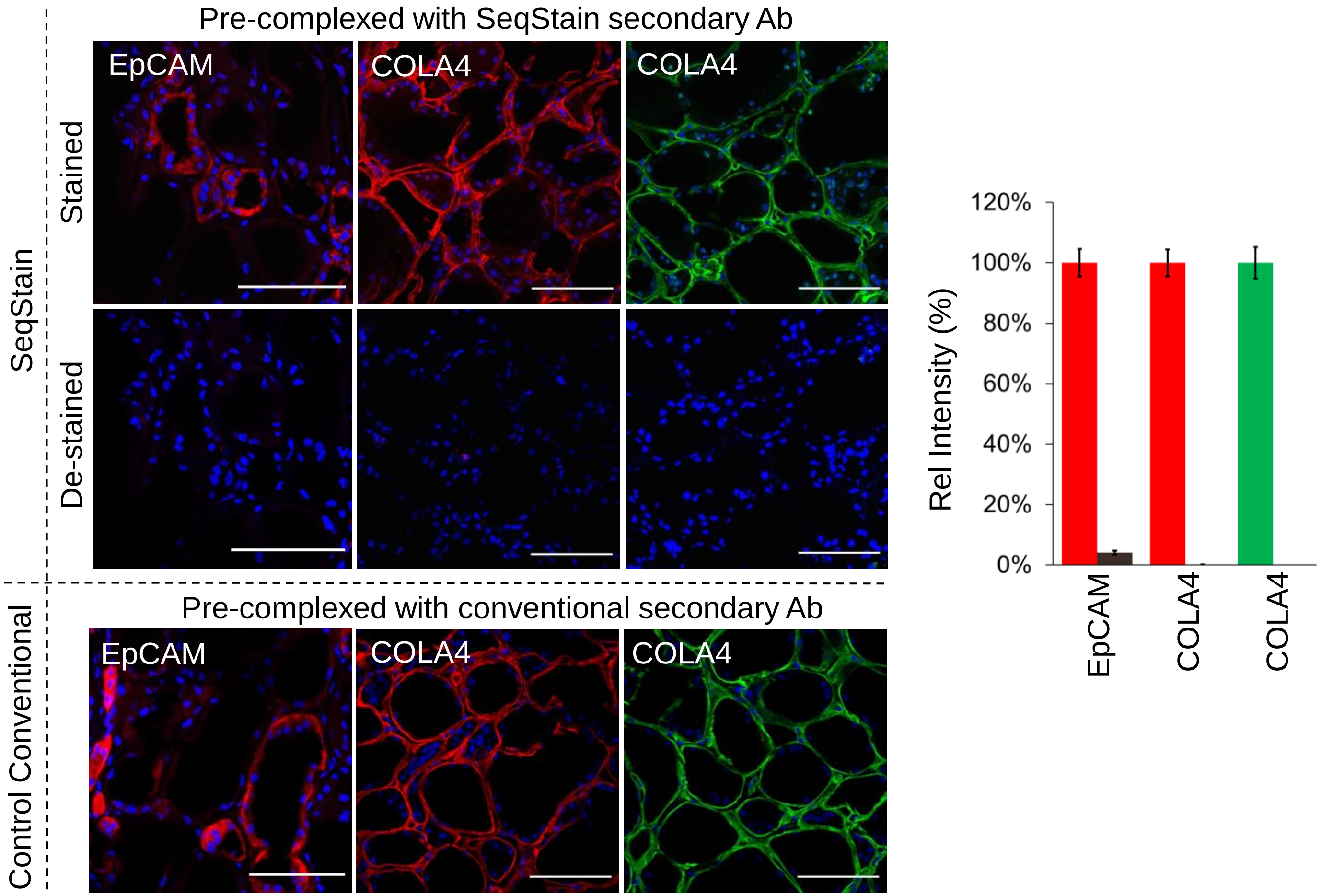
Pre-complexed with SeqStain secondary Ab
EpCAM
COLA4
COLA4
SeqStain
Stained
De-stained
Pre-complexed with conventional secondary Ab
Control Conventional
EpCAM
COLA4
COLA4
Rel Intensity (%)
120%
100%
80%
60%
40%
20%
0%
EpCAM
COLA4
COLA4

Figure S12

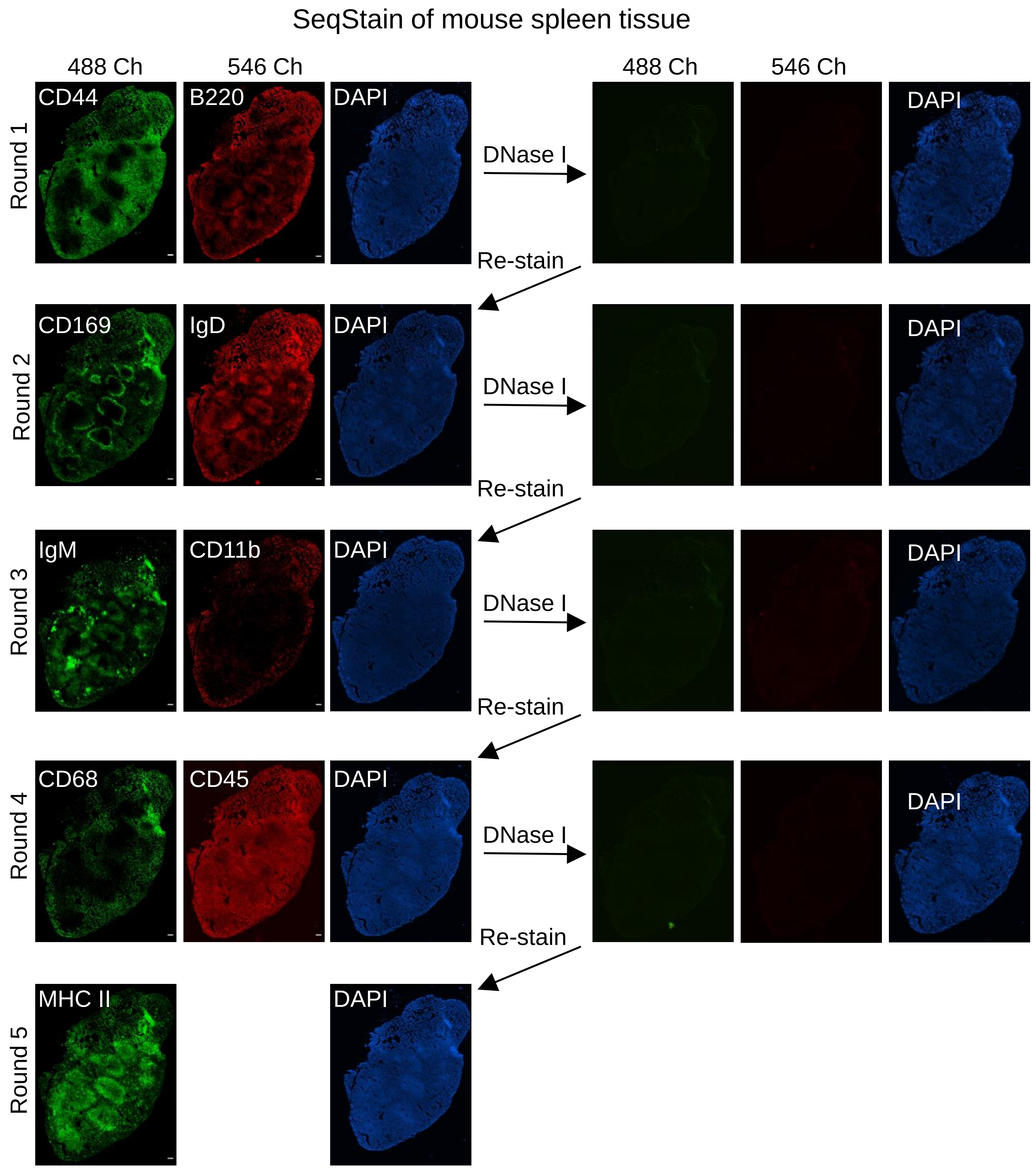
SeqStain of mouse spleen tissue
488 Ch
546 Ch
488 Ch
546 Ch
Round 1
CD44
B220
DAPI
DNase I
DAPI
Re-stain
Round 2
CD169
IgD
DAPI
DNase I
DAPI
Re-stain
Round 3
IgM
CD11b
DAPI
DNase I
DAPI
Re-stain
Round 4
CD68
CD45
DAPI
DNase I
DAPI
Re-stain
Round 5
MHC II
DAPI

Figure S13

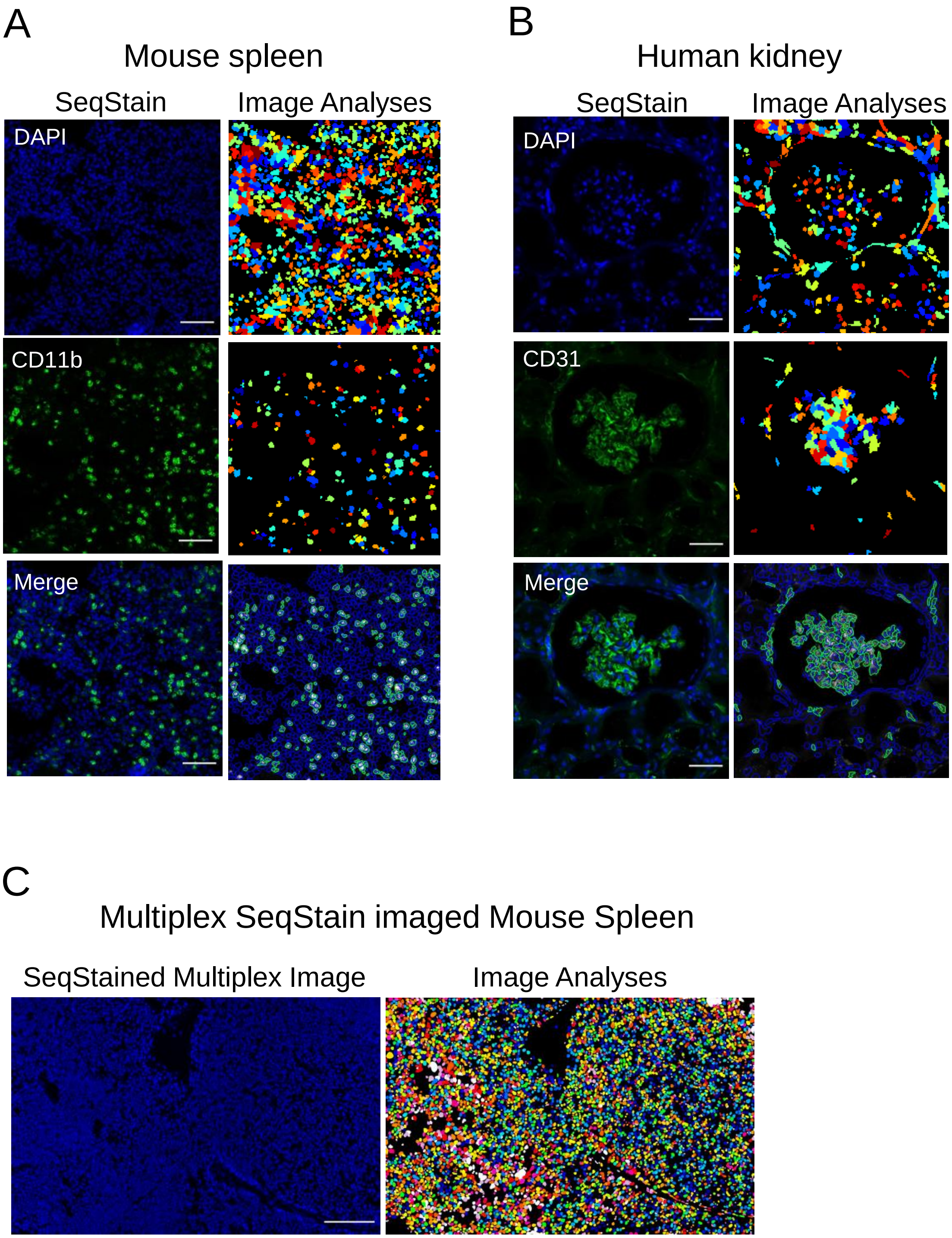
A
Mouse spleen
SeqStain
Image Analyses
DAPI
CD11b
Merge
B
Human kidney
SeqStain
Image Analyses
DAPI
CD31
Merge
C
Multiplex SeqStain imaged Mouse Spleen
SeqStained Multiplex Image
Image Analyses

Figure S14

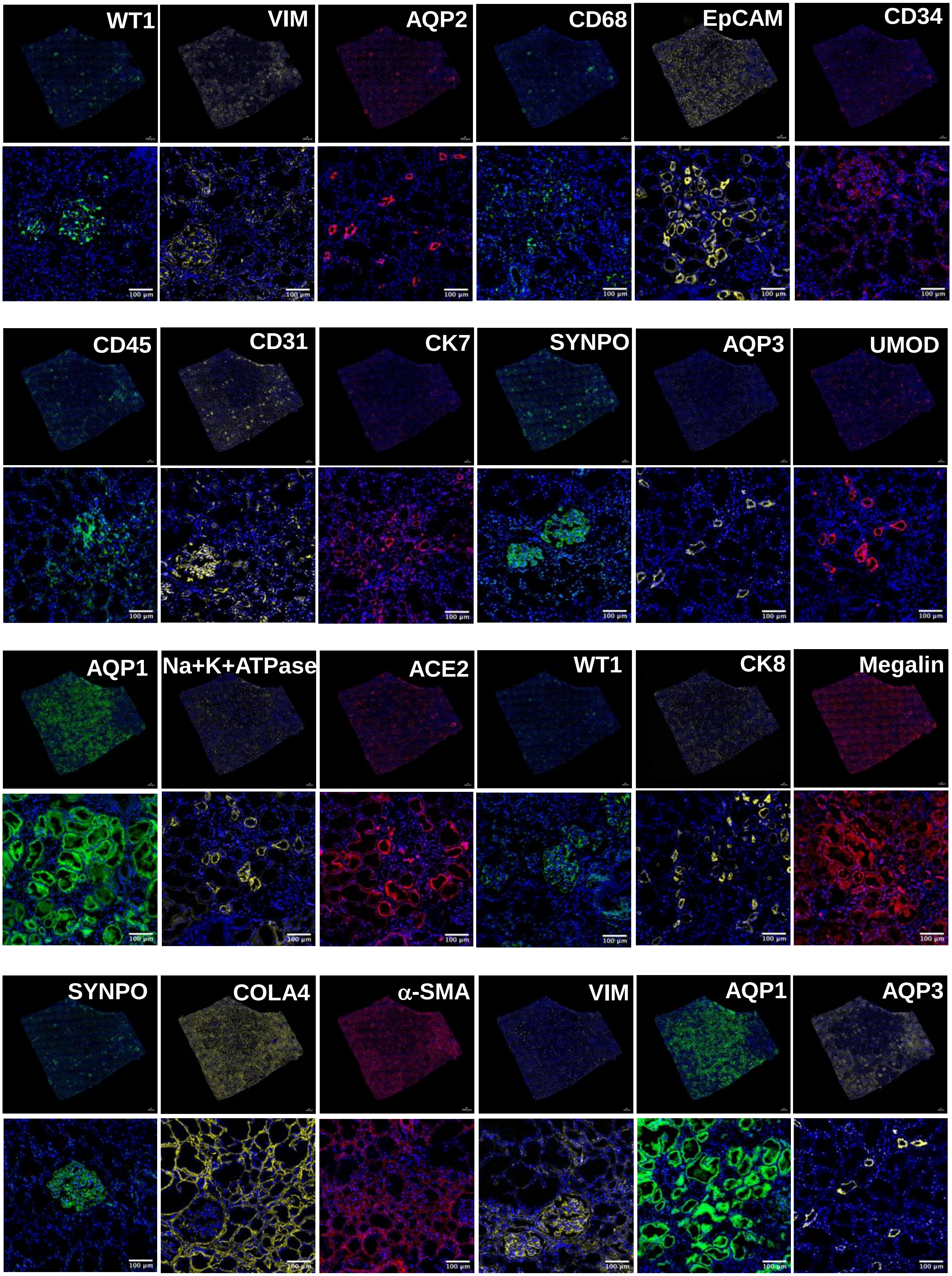
WT1
VIM
AQP2
CD68
EpCAM
CD34
CD45
CD31
CK7
SYNPO
AQP3
UMOD
AQP1
Na+K+ATPase
ACE2
WT1
CK8
Megalin
SYNPO
COLA4
α-SMA
VIM
AQP1
AQP3
100 μm

Figure S15

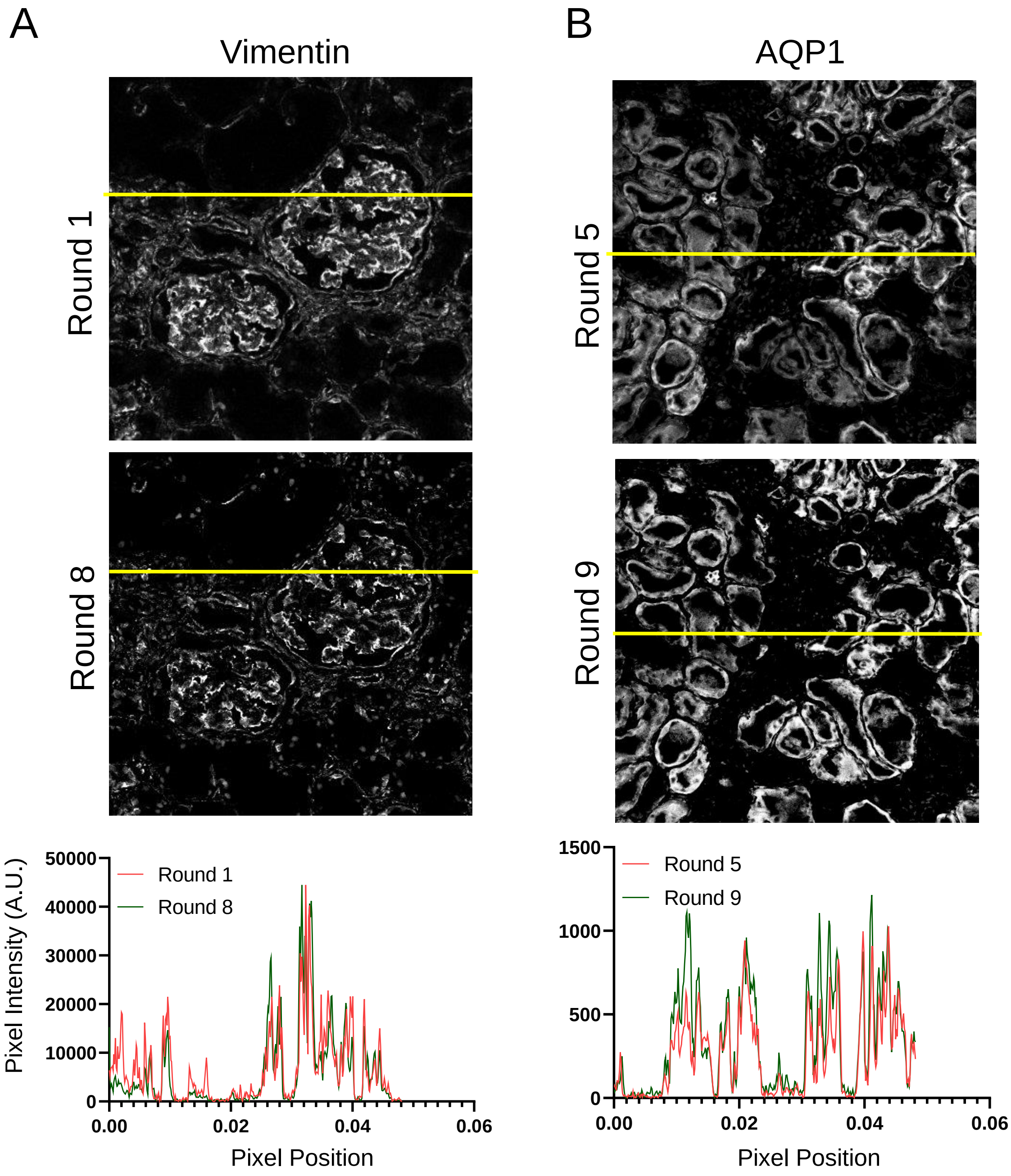
A
Vimentin
Round 1
Round 8
B
AQP1
Round 5
Round 9
Pixel Intensity (A.U.)
50000
40000
30000
20000
10000
0
Round 1
Round 8
0.00
0.02
0.04
0.06
Pixel Position
1500
1000
500
0
Round 5
Round 9
0.00
0.02
0.04
0.06
Pixel Position

Figure S16
